# Rare variant analysis of 4,241 pulmonary arterial hypertension cases from an international consortium implicate *FBLN2*, *PDGFD* and rare *de novo* variants in PAH

**DOI:** 10.1101/2020.05.29.124255

**Authors:** Na Zhu, Emilia M. Swietlik, Carrie L. Welch, Michael W. Pauciulo, Jacob J. Hagen, Xueya Zhou, Yicheng Guo, Johannes Karten, Divya Pandya, Tobias Tilly, Katie A. Lutz, Erika Rosenzweig, Usha Krishnan, Anna W. Coleman, Claudia Gonzaga-Juaregiu, Allan Lawrie, Richard C. Trembath, Martin R. Wilkins, Regeneron Genetics Center, PAH Biobank Enrolling Centers’ Investigators, NIHR BioResource for Translational Research - Rare Diseases, National Cohort Study of Idiopathic and Heritable PAH, Nicholas W. Morrell, Yufeng Shen, Stefan Gräf, William C. Nichols, Wendy K. Chung

## Abstract

**Background:** Group 1 pulmonary arterial hypertension (PAH) is a lethal vasculopathy characterized by pathogenic remodeling of pulmonary arterioles leading to increased pulmonary pressures, right ventricular hypertrophy and heart failure. Recent high-throughput sequencing studies have identified additional PAH risk genes and suggested differences in genetic causes by age of onset. However, known risk genes explain only 15-20% of non-familial idiopathic PAH cases.

**Methods:** To identify new risk genes, we utilized an international consortium of 4,241 PAH cases with 4,175 sequenced exomes (n=2,572 National Biological Sample and Data Repository for PAH; n=469 Columbia University Irving Medical Center, enriched for pediatric trios) and 1,134 sequenced genomes (UK NIHR Bioresource – Rare Diseases Study). Most of the cases were adult-onset disease (93%), and 55% idiopathic (IPAH) and 35% associated with other diseases (APAH). We identified protein-coding variants and performed rare variant association analyses in unrelated participants of European ancestry, including 2,789 cases and 18,819 controls (11,101 unaffected parents from the Simons Powering Autism Research for Knowledge study and 7,718 gnomAD individuals). We analyzed *de novo* variants in 124 pediatric trios.

**Results:** Seven genes with rare deleterious variants were significantly associated (false discovery rate <0.1) with IPAH, including three known genes (*BMPR2*, *GDF2*, and *TBX4*), two recently identified candidate genes (*SOX17*, *KDR*), and two new candidate genes (*FBLN2*, fibulin 2; *PDGFD*, platelet-derived growth factor D). The candidate genes exhibit expression patterns in lung and heart similar to that of known PAH risk genes, and most of the variants occur in conserved protein domains. Variants in known PAH gene, *ACVRL1*, showed association with APAH. Predicted deleterious *de novo* variants in pediatric cases exhibited a significant burden compared to the background mutation rate (2.5x, p=7.0E-6). At least eight novel candidate genes carrying *de novo* variants have plausible roles in lung/heart development.

**Conclusions:** Rare variant analysis of a large international consortium identifies two new candidate genes - *FBLN2* and *PDGFD*. The new genes have known functions in vasculogenesis and remodeling but have not been previously implicated in PAH. Trio analysis predicts that ~15% of pediatric IPAH may be explained by *de novo* variants.

## Background

Pulmonary arterial hypertension (PAH) remains a progressive, lethal vasculopathy despite recent therapeutic advances. The disease is characterized by pulmonary vascular endothelial dysfunction and proliferative remodeling giving rise to increased pulmonary pressures and resistance, heart failure and high mortality (1–3). Dysregulated vascular, inflammatory and immune cells contribute to these pathological processes (3). PAH can present at any age, but the ~3:1 female to male ratio in adult-onset disease is not observed in pediatric-onset disease, in which the disease incidence is similar for males and females. The estimated prevalence of PAH is 4.8-8.1 cases/million for pediatric-onset (4) and 5.6-25 cases/million for adult-onset disease (5). Early genetic linkage and candidate gene studies indicated an autosomal dominantly-inherited genetic component to risk for disease. However, the susceptibility variants are incompletely penetrant, many individuals who carry monogenic risk variants never develop PAH, and a subset of patients have deleterious variants in more than one risk gene. These data suggest that additional genetic, epigenetic, environmental factors, and gene x environment interactions, contribute to disease.

Genetic analyses of larger cohorts using gene panels, exome sequencing (ES) or genome sequencing (GS) have further defined the frequency of individuals with deleterious variants in PAH risk genes and have identified novel candidate risk genes. Bone morphogenetic protein receptor type 2 (*BMPR2*) mutations are observed in 60-80% of familial (FPAH) cases across genetic ancestries (6–10). *BMPR2* carriers have younger mean age-of-onset and are less responsive to vasodilators compared to non-carriers(6, 11, 12), with an enrichment of predicted deleterious missense (D-Mis) variants with younger age-of-onset (6, 13). Notably, *BMPR2* variants contribute almost exclusively to FPAH and idiopathic PAH (IPAH), and rarely to other subclasses. Variants in two other genes in the transforming growth factor-beta (TGF-β) superfamily, activin A receptor type II-like 1 (*ACVRL1*) and endoglin (*ENG*) contribute to ~0.8% of PAH cases(6), especially PAH associated with hereditary hemorrhagic telangiectasia (APAH-HHT). Variants in growth differentiation factor 2 (*GDF2*), encoding the ligand of BMPR2/ACVRL1 (BMP9), contribute to ~1% of PAH cases in European-enriched cohorts (6, 7) and are more frequent in Chinese patients (~6.7%)(14). Variants in mothers against decapentaplegic (*SMAD*) genes, encoding downstream mediators of BMP signaling, contribute rarely.

A number of genes outside of the TGF-β signaling pathway have also been identified as PAH risk genes. Variants in developmental transcription factors*, TBX4* and *SOX17*, are enriched in pediatric patients(6, 15–17). Variants in *TBX4* cause a variety of developmental lung disorders (18), including persistent pulmonary hypertension of the newborn with recurrent PAH in childhood (19). Variants in *SOX17* contribute to ~3.2% of PAH associated with congenital heart disease (APAH-CHD)(17). Biallelic variants in eukaryotic initiation translation factor (*EIF2AK4*) cause pulmonary veno-occlusive disease (PVOD) and pulmonary capillary hemangiomatosis (PCH)(20, 21). Loss of function variants in channelopathy genes potassium two pore domain channel (*KCNK3*)(22) and ATP-binding cassette subfamily member 8 (*ABCC8*)(23), as well as membrane reservoir gene caveolin-1 (*CAV1*)(24–26), are causative for PAH. Recent associations of variants in ATPase 13A3 (*ATP13A3*) and aquaporin 1 (*AQP1*)(7), as well as kallikrein 1 (*KLK1*) and gamma-glutamyl carboxylase (*GGCX*)(6) have been recently reported but require independent confirmation. Finally, a role for *de novo* variants in pediatric-onset PAH has been suggested based on a cohort of 34 child-parent trios(16).

Together, these data indicate that rare genetic variants underlie ~75-80% of FPAH(27), at least 10% of adult-onset idiopathic PAH (IPAH)(6, 7) and up to ~36% of pediatric-onset IPAH(28). A substantial fraction of non-familial PAH cases remains genetically undefined. The low frequency of risk variants for each gene, except *BMPR2*, indicates that large numbers of individuals are required for further validation of rare risk genes and pathways, and to understand the natural history of each genetic subtype of PAH. Towards this end, we analyzed 4,175 PAH cases from an international consortium with ES or GS. In addition, we analyzed *de novo* mutations in an expanded cohort of 124 pediatric trios.

## METHODS

### Patient cohorts and control datasets

A total of 4,175 PAH cases from the National Biological Sample and Data Repository for PAH (PAH Biobank, n=2,570 exomes)(6), UK NIHR Bioresource – Rare Diseases Study (UK NIHR Bioresource, n=1,144 genomes)(7) and the Columbia University Irving Medical Center (CUIMC, n=461 exomes)(16, 17, 23) were included in a combined analysis of rare inherited variants. The subset of 124 affected child-unaffected parents trios (n=111 CUIMC, n=8 UK NIHR Bioresource, n=5 PAH Biobank) were included in an analysis of *de novo* variants. An additional 65 *BMPR2* mutation-positive cases from CUIMC without exome sequencing data were previously-reported (16, 17) and included in the overall cohort counts (total of 4,241 cases). As previously described, cases were diagnosed by medical record review including right heart catheterization and all were classified as World Symposium on Pulmonary Hypertension (WSPH) PH group I (29). Written informed consent for publication was obtained at enrollment. The studies were approved by the institutional review boards at CCHMC, individual PAH Biobank Centers, the East of England Cambridge South national research ethics committee (REC, ref. 13/EE0325) or CUIMC.

The control group consisted of unaffected parents from the Simons Powering Autism Research for Knowledge (SPARK) study (exomes) (30) as well as gnomADv2.1.1 (gnomAD) individuals (genomes).

### ES/GS data analysis

PAH Biobank, CUIMC and SPARK cohort samples were all sequenced in collaboration with the Regeneron Genetics Center as previously described (6, 7, 16, 17, 23); the UK NIHR Bioresource sequence data were also previously described (7). For case and SPARK control data, we used a previously established bioinformatics procedure (31) to process and analyze exome and genome sequence data. For the UK NIHR Bioresource data, we extracted reads from GS data by the following procedure: 1) obtained all erads that were mapped to the human genome regions that overlapped with the target regions of xGEN exome capture intervals (Exome Research panel 1.0); 2) the mate pairs of these reads. We then processed the extracted GS data using the same pipeline as the ES data. Specifically, we used BWA-MEM(32) to map and align paired-end reads to the human reference genome (version GRCh38/hg38, accession GCA 000001405.15), Picard v1.93 MarkDuplicates to identify and flag PCR duplicates and GATK v4.1(33, 34) HaplotypeCaller in Reference Confidence Model mode to generate individual-level gVCF files from the aligned sequence data. We then performed joint calling of variants from all three datasets using *GLnexus* (35). We used the following inclusion rules to select variants for downstream analysis: AF<0.05% in the cohort, <0.01% in gnomAD exome_ALL (all ancestries), >90% target region with dp>=10; mappability=1; allele balance>=0.25; and “PASS” from DeepVariants(36). For gnomAD data, only variants located in xGen-captured protein coding regions were used, and filtering was based on GATK metrics obtained from gnomAD. For cases and SPARK controls, we called variants from each sample using DeepVariants, a new tool based on machine learning (37). We used the ES mode for ES data and GS mode for GS data. Variants used for downstream analyses were restricted to the subset called by both *GLnexus* and DeepVariants.

*De novo* variants were defined as a variant present in the offspring with homozygous reference genotypes in both parents. We used a series of filters to identify *de novo* variants: VQSR tranche ≤99.7 for SNVs and ≤99.0 for indels; GATK Fisher Strand ≤25; quality by depth ≥2. We required the candidate *de novo* variants in probands to have ≥5 reads supporting the alternative allele, ≥20% alternative allele fraction, Phred-scaled genotype likelihood ≥60 (GQ), and population AF ≤0.01% in ExAC; and required both parents to have ≥10 reference reads, <5% alternative allele fraction, and GQ ≥30.

We used Ensembl Variant Effect Predictor (VEP, Ensemble 93)(38) to annotate variant function and ANNOVAR(39) to aggregate variant population frequencies and *in silico* predictions of deleteriousness. Rare variants were defined as AF ≤0.01% in gnomAD exome_ALL (all ancestries). Deleterious variants were defined as likely gene-disrupting (LGD, including premature stop-gain, frameshift indels, canonical splicing variants and exon deletions) or predicted damaging missense (D-Mis) based on gene-specific REVEL score thresholds(17, 40) (see below). All inherited and *de novo* rare variants in candidate genes were manually inspected using Integrative Genome Viewer (IGV).

### Statistical analysis

To identify novel risk genes for IPAH, we performed a rare variant association test in unrelated participants of European ancestry. Genetic ancestry and relatedness of cases and SPARK controls were checked using Peddy (41), and only unrelated cases (n = 2,789) and controls (18,819: 11,101 SPARK parents and 7,718 gnomAD individuals) were included in the association test. The gnomAD controls were confined to non-Finish Europeans (NFE). We performed a gene-based case-control test comparing the frequency of rare deleterious variants in PAH cases with unaffected controls. To reduce batch effects in combined datasets from different sources (42), we limited the analysis to regions targeted by xGen and with at least 10X coverage in 90% of samples. We then tested for similarity of the rare synonymous variant rate among cases and controls, assuming that most rare synonymous variants do not have discernible effects on disease risk.

To identify PAH risk genes, we tested the burden of rare deleterious variants (AF ≤0.01%, LGD or D-Mis) in each protein-coding gene in cases compared to controls using a variable threshold test. Specifically, we used REVEL(40) scores to predict the deleteriousness of missense variants, searched for a gene-specific optimal REVEL score threshold that maximized the burden of rare deleterious variants in cases compared to controls, and then used permutations to calculate statistical significance as described previously(6) to control type I error rate (6). We checked for inflation using a quantile-quantile (Q-Q) plot and calculated the genomic control factor, lambda, using QQperm (https://cran.r-project.org/web/packages/QQperm/QQperm.pdf). Lambda equal to 1 indicates no deviation from the expected distribution. We performed two association tests, one with LGD and D-Mis variants combined and the other with D-Mis variants alone. We defined the threshold for genome-wide significance by Bonferroni correction for multiple testing (n=40,000, as two tests for each gene and there are 18,939 protein-coding genes, threshold p-value=1.25e-6). We used the Benjamini-Hochberg procedure to estimate false discovery rate (FDR) by p.adjust in R.

To estimate the burden of *de novo* variants in cases, we calculated the background mutation rate using a previously published tri-nucleotide change table(43, 44) and calculated the rate in protein-coding regions that are uniquely mappable. We assumed that the number of *de novo* variants of various types (e.g. synonymous, missense, LGD) expected by chance in gene sets or all genes followed a Poisson distribution(43). For a given type of *de novo* variant in a gene set, we set the observed number of cases to *m1*, the expected number to *m0*, estimated the enrichment rate by (*m1*/*m0*) and tested for significance using an exact Poisson test (poisson.test in R) with *m0* as the expectation.

### Protein modeling

Homology structures of conserved protein domains in FBLN2 and PDGFD were built using EasyModeller 4.0(45). Template structures were downloaded from the protein database (PDB) for endothelial growth factor (EGF, PDB ID 5UK5) and CUB (PDB ID 3KQ4) domains. The template structure for platelet-derived growth factor (PDGF)/vascular EGF (VEGF) was downloaded directly from PrePPI(46, 47).

### Gene expression

Single cell RNA-seq data of aorta, lung, and heart tissues were obtained from *Tabula Muris*, a transcriptome compendium containing RNA-seq data from ~100,000 single cells from 20 adult-staged mouse organs (48). We chose 14 tissue/cell types including endothelial, cardiac muscle, and stromal cells from the three tissues, restricting the analysis to tissues/cell types for which there was RNA-seq data from at least 70 individual cells (Supplementary Figure 5a). Relative gene expression was based on the fraction of cells with >0 reads in each cell type. PCA of cell type specific gene expression profiles was performed and the script is available through GitHub (https://github.com/ShenLab/US_UK_combine).

## RESULTS

### Cohort characteristics

Demographic data and mean hemodynamic parameters of the combined US/UK cohort are shown in Table 1. The cohort includes 4,241 cases: 54.6% IPAH, 34.8% APAH, 5.9% FPAH and 4.6% other PAH. Most of the APAH and other PAH cases came from the PAH Biobank and have been described previously(6). The majority of cases were female (75.1%) and adult-onset (92.6%) with a mean age-of-diagnosis (by right heart catheterization) of 45.9 ± 20 years (mean ± SD). The genetically-determined ancestries were European (74.5%), Hispanic (8.6%), African (8.7%), East Asian (2.5%), and South Asian (2.8%). Hemodynamic data were collected at the time of PAH diagnosis. The mean pulmonary arterial pressure (mPAP) and mean pulmonary capillary wedge pressure (mPCWP) for the overall cohort was 51 ± 14 mm Hg (mean ± SD) and 10 ± 4 mm Hg, respectively, compared to 58 ± 14 mm Hg and 10 ± 4 mm Hg for FPAH.

**Table 1.** Demographic data and mean hemodynamic parameters from the US/UK PAH cohort*.

|  | **All** | **IPAH** | **APAH\*\*** | **FPAH** | **Other\*\*\*** |
| --- | --- | --- | --- | --- | --- |
| **Total, n (%)** | 4241 | 2319 (54.6) | 1479 (34.8) | 252 (5.9) | 191 (4.6) |
| **Sex, n (%)** |  |  |  |  |  |
| F | 3187 (75.1) | 1721 (74.2) | 1156 (78.2) | 173 (68.6) | 137 (71.7) |
| M | 1054 (24.9) | 598 (25.8) | 323 (21.8) | 79 (31.3) | 54 (28.3) |
| F:M ratio | 3:1 | 2.9:1 | 3.6:1 | 2.3:1 | 2.5:1 |
| **Age-of-onset, n (%)** |  |  |  |  |  |
| Adult ($\geq 18$ y) | 3780 (89.2) | 2126 (91.7) | 1256 (85.2) | 213 (84.5) | 185 (96.9) |
| Child ($< 18$ y) | 457 (10.8) | 193 (8.3) | 219 (14.8) | 39 (15.4) | 6 (3.1) |
| Mean ± SD | 45.9 ± 20.0 | 47.0 ± 19.5 | 45.2 ± 21.3 | 36.8 ± 16.8 | 47.7 ± 15.0 |
| **Ancestry, n (%)** |  |  |  |  |  |
| EUR | 3108 (74.5) | 1798 (77.5) | 988 (67.0) | 166 (86.9) | 156 (81.7) |
| HISP | 359 (8.6) | 145 (6.3) | 186 (12.6) | 13 (6.8) | 15 (7.9) |
| AFR | 365 (8.7) | 166 (7.2) | 189 (12.8) | 2 (1.0) | 8 (4.2) |
| EAS | 104 (2.5) | 42 (1.8) | 57 (3.9) | 1 (0.5) | 4 (2.1) |
| SAS | 116 (2.8) | 82 (3.5) | 28 (1.9) | 4 (2.1) | 2 (1.0) |
| Other/unknown | 123 (2.9) | 85 (3.7) | 27 (1.8) | 5 (2.6) | 6 (3.1) |
| **Hemodynamic parameters, mean ± SD (n)** |  |  |  |  |  |
| MPAP, mmHg | 51 ± 14 (3594) | 53 ± 17 (2045) | 48 ± 14 (1235) | 58 ± 14 (158) | 51 ± 12 (156) |
| MPCWP, mmHg | 10 ± 4 (3407) | 10 ± 24 (1912) | 10 ± 4 (1197) | 10 ± 4 (151) | 11 ± 4 (147) |

\*US/UK PAH cohort: 2,572 PAH Biobank; 1,134 UK NIHR Bioresource; 534 CUIMC.
\*\*APAH: PAH associated with connective tissue diseases, congenital heart disease, HHT, HIV.
\*\*\*Other: diet- and toxin-induced PAH, non-familial pulmonary veno-occlusive disease/pulmonary capillary hemangiomatosis and one case of persistent pulmonary hypertension of the newborn.
Abbreviations: AFR, African; EAS, East Asian; EUR, European; HISP, Hispanic; SAS, Southeast Asian; MPAP, mean pulmonary artery pressure; MPCWP, mean pulmonary capillary wedge pressure.

A comparison of the clinical characteristics and hemodynamic data for pediatric-versus adult-onset PAH cases is shown in Supplementary Table 1. Notably, the female:male ratio among pediatric-onset cases was significantly lower (1.65:1) compared to adult-onset cases (4:1, p <0.0001 by Fisher’s exact test), and children had higher mPAP and mPCWP, decreased cardiac output and increased pulmonary vascular resistance compared to adults at diagnosis (all differences p <0.0001 by Student’s t-test).

Rare deleterious variants in *BMPR2* were identified in 7.7% of cases overall (209/2318, 9% of IPAH, 108/191, 56.6% of FPAH, and 13/1475, 0.88% of APAH). The variants include LGD and D-Mis variants as well as intragenic or whole gene deletions as previously described (6, 7, 16, 17). The percentage of *BMPR2* carriers in the US/UK international cohort is lower than previous reports (7, 11) due to the enrichment of APAH cases, rarely caused by *BMPR2* variants (6, 17).

### Identification of novel risk genes: *FBLN2* and *PDGFD*

To perform a combined analysis of US and UK sequencing data, we reprocessed the UK data using our inhouse pipeline, including predictions of missense variant deleteriousness(6). Quality control procedures included detection of cryptic relatedness amongst all PAH participants. We performed a gene-based case-control association analysis to identify novel PAH risk genes using only unrelated cases. To control for population stratification, we confined the association analysis to individuals of European ancestry (2,789 cases, 18,819 controls) and then screened the whole cohort, including nonEuropeans, for rare deleterious variants in associated genes. As a quality control check for the filtering parameters employed, we compared the frequencies of rare synonymous variants, a variant class that is mostly neutral with respect to disease status, in European cases vs controls. We observed similar frequencies of synonymous variants in cases vs controls (enrichment rate =1.0, p-value =0.28) (Supplementary Table 2). Furthermore, a gene-level burden test revealed no enrichment of rare synonymous variants in cases (Supplementary Figure 1). We then proceeded to test for gene-specific enrichment of rare deleterious variants (AF <0.01%, LGD and D-Mis, or D-Mis only) in cases compared to controls. To improve power, we empirically determined the optimal REVEL score threshold to define deleterious missense variants in a gene-specific manner using a variable threshold test(6). To account for potential different modes of action for different risk genes, we tested the association twice for each gene: one with LGD and D-Mis variants and the other with D-Mis variants alone. In this approach, LGD and D-Mis together is optimized for complete or partial loss of function; D-Mis alone is optimized for gain of function or dominant negative variants. We set the total number of tests at twice the number of protein-coding genes for multiple test adjustment, a conservative approach considering that the data used in these two tests per gene are not independent. The Q-Q plot of p-values from tests in all genes shows negligible genomic inflation (Supplementary Figure 2). Rare deleterious variants in eleven genes were significantly associated (false discovery rate, FDR <0.1) with PAH. Among these, seven are known or previously-reported candidate PAH risk genes: *BMPR2*, *TBX4*, *GDF2*, *ACVRL1*, *SOX17*, *AQP1*, *ATP13A3*, and *KDR*. Three are new candidate genes: *COL6A5* (collagen type VI alpha 5 chain), *JPT2* (Jupiter microtubule-associated homolog 2) and *FBLN2* (fibulin 2).

To take advantage of the relatively large number of European IPAH cases in the combined cohort, we then restricted the analysis to IPAH. Again, testing for association across all protein-coding genes for 1,647 IPAH cases compared to 18,819 controls was generally consistent with expectation under the null model (Figure 1). Rare predicted deleterious variants in seven genes were significantly associated (FDR <0.1) with IPAH, including three known genes (*BMPR2*, *GDF2*, and *TBX4*), two recently identified candidate genes (*SOX17* and *KDR*) and two new candidate genes (*FBLN2*, and *PDGFD*, platelet-derived growth factor D). More than 95% of samples for both cases and controls had at least 10X depth of sequence coverage across the target regions for *FBLN2* and *PDGFD* (Supplementary Figure 3), excluding the possibility that the associations were driven by coverage differences between cases and controls. We also tested for gene-level associations restricting the analysis to European APAH cases (n=998). The Q-Q plot of p-values from all gene tests is shown in Supplementary Figure 4. Known PAH gene *ACVRL1* showed association with APAH, consistent with its role in APAH-HHT, but no genes were significantly associated at FDR <0.1.

**Figure 1.**
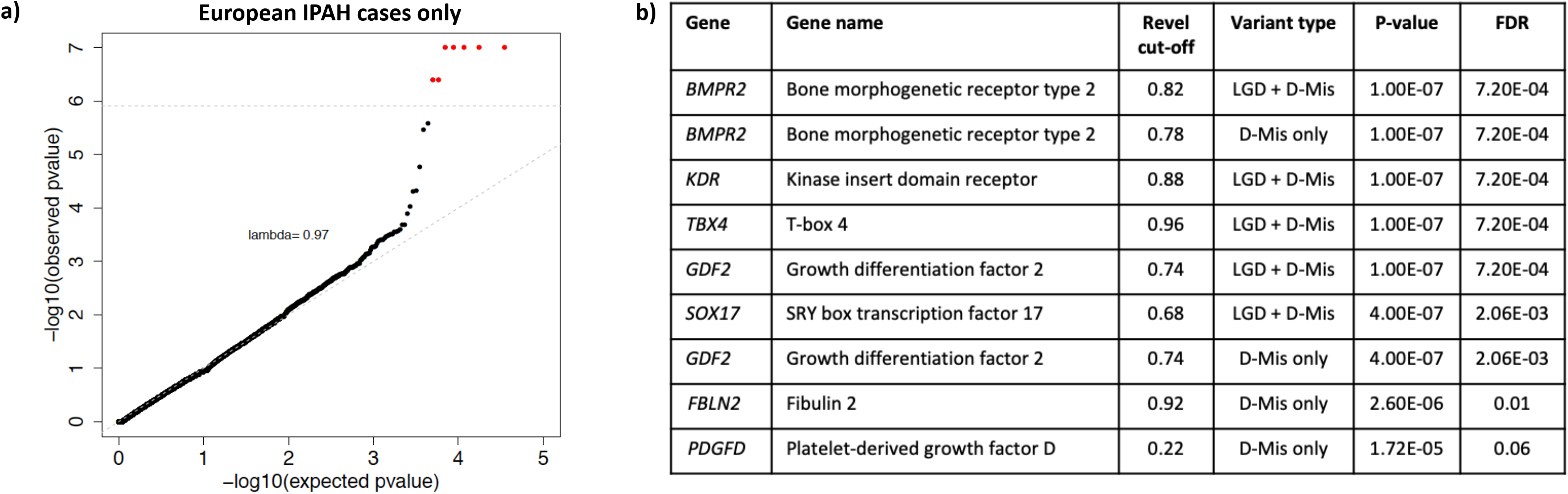
Gene-based association analysis using 1,647 European IPAH cases and 18,819 European controls. Controls include 11,101 SPARK parents and 7,718 NFE gnomAD v2.1.1. **a)** Results of a binomial test confined to rare, likely gene damaging (LGD) and predicted deleterious missense (D-Mis) variants or D-Mis only variants in 20,000 protein-coding genes. Horizontal gray line indicates the Bonferroni-corrected threshold for significance. **b)** Complete list of top association genes (FDR<0.1).

*KDR* has recently been implicated as a causal gene for PAH based on a small familial study (49) and our population-based phenotype-driven (SKAT-O) analysis of the UK NIHR Bioresource cohort with replication in the PAH Biobank (50). Both of those analyses were based on protein truncating variants. Herein, we provide additional statistical evidence based on a burden test including both LGD and D-Mis variants using our variable threshold method. Six cases (5 IPAH, 1APAH-CHD) carry D-Mis variants with empirically-determined REVEL >0.86; none of these cases have variants in other known PAH risk genes. One of the variants, c.3439C>T is recurrent in three cases. The age-of-onset for the six cases is 57 ± 20 years (mean±SD, range 25-75 years); all are of European ancestry. Details of the variants are provided in Supplementary Table 3. All of the variants are located in the conserved tyrosine kinase domain of the encoded protein (www.uniprot.org). Statistically-significant association following Bonferroni correction for multiple testing provides confirmation of the association of *KDR* with PAH using an alternative burden-based statistical method.

The associations of *FBLN2* and *PGDFD* were both driven by D-Mis variants. We next screened the entire combined cohort, including participants of non-European ancestry, for rare deleterious missense variants in *FBLN2* and *PDGFD*. In total, seven cases carry *FBLN2* variants (6 IPAH, 1 APAH) and ten cases carry *PDGFD* variants (9 IPAH, 1 PAH associated with diet and toxins) (Table 2). Most of the carriers are of European ancestry; one *FBLN2* carrier is of East Asian ancestry and one *PDGFD* carrier is of African ancestry. One *FBLN2* variant ((c.2944G>T; p.(Asp982Tyr)) and two *PDGFD* variants ((c.385G>A; p.(Glu129Lys) and c.961T>A; p.(Tyr321Asn)) were recurrent in the cohort. Locations of the predicted damaging missense amino acid residues are shown in Figure 2. FBLN2 contains multiple endothelial growth factor (EGF) domains, and PDGFD contains a conserved CUB domain and a platelet-derived growth factor (PDGF)/vascular EGF (VEGF) domain. All of the *FBLN2* and eight out of ten *PDGFD* D-Mis variants, occur in conserved protein domains. FBLN2 p.(Gly880Val) and p.(Gly889Asp) replace conserved reverse turn residues in an EGF domain which may change the conformation of the domain and impact protein function (Figure 2b). Recurrent FBLN2 p.(Asp982Tyr) disrupts a Ca^++^ binding site(51) in another EGF domain (Figure 2b), which may reduce the affinity and frequency of Ca^++^ binding. PDGFD p.(Asp148Asn) disrupts a Ca^++^ binding site within the CUB domain(52) (Figure 2c) and recurrent PDFGD p.(Tyr321Asn) is predicted to disrupt a hydrogen bond within the PDGF/VEGF domain (Figure 2c). In addition, PDGFD p.(Arg295Cys) is located in close proximity to Cys356 and Cys358, potentially introducing new disulfide bonds within the PDGF/VEGF domain.

**Table 2.** Rare predicted deleterious *FBLN2* and *PDGFD* variants* among 4,175 PAH cases**.

| Case ID | Sex | Age<sub>dx</sub> | PAH subclass | Ancestry | Gene *** | Exon | Nucleotide change | Amino acid change | Variant type | MAF (gnomAD exomes) | CADD score | Revel |
| --- | --- | --- | --- | --- | --- | --- | --- | --- | --- | --- | --- | --- |
| 08-018 | F | 70 | IPAH | EUR | *FBLN2* | 12 | c.2639G>T | p.(Gly880Val) | D-Mis | 1.63E-05 | 27.1 | 0.94 |
| 17-035 | F | 41 | APAH | EAS | *FBLN2* | 12 | c.2666G>A | p.(Gly889Asp) | D-Mis | --- | 27.6 | 0.94 |
| 12-207 | F | 44 | IPAH | EUR | *FBLN2* | 13 | c.2794T>C | p.(Phe923Leu) | D-Mis | 2.11E-05 | 29.6 | 0.92 |
| 23-001 | M | 66 | IPAH | EUR | *FBLN2* | 14 | c.2944G>T | p.(Asp982Tyr) | D-Mis | 1.88E-05 | 34.0 | 0.95 |
| 29-031 | F | 57 | IPAH | EUR | *FBLN2* | 14 | c.2944G>T | p.(Asp982Tyr) | D-Mis | 1.88E-05 | 34.0 | 0.95 |
| 34-005 | M | 69 | IPAH | EUR | *FBLN2* | 14 | c.2944G>T | p.(Asp982Tyr) | D-Mis | 1.88E-05 | 34.0 | 0.95 |
| W000210 | F | 52 | IPAH | EUR | *FBLN2* | 14 | c.2944G>T | p.(Asp982Tyr) | D-Mis | 1.88E-05 | 34.0 | 0.95 |
| W000073 | F | 40 | IPAH | AFR | *PDGFD* | 2 | c.166G>A | p.(Gly56Ser) | D-Mis | 1.99E-05 | 22.9 | 0.64 |
| JM950 | F | 2 | IPAH | EUR | *PDGFD* | 2 | c.250C>T | p.(Arg84Trp) | D-Mis | 1.59E-05 | 16.4 | 0.51 |
| E012465 | F | 55 | IPAH | EUR | *PDGFD* | 3 | c.385G>A | p.(Glu129Lys) | D-Mis | --- | 25.2 | 0.262 |
| E014342 | F | 40 | IPAH | EUR | *PDGFD* | 3 | c.385G>A | p.(Glu129Lys) | D-Mis | --- | 25.2 | 0.262 |
| E014400 | F | 43 | IPAH | EUR | *PDGFD* | 3 | c.442G>A | p.(Asp148Asn) | D-Mis | 7.97E-06 | 25.2 | 0.41 |
| E000844 | F | 39 | IPAH | EUR | *PDGFD* | 6 | c.770T>C | p.(Leu257Pro) | D-Mis | 4.01E-06 | 31.0 | 0.62 |
| 13-037 | M | 43 | DTOX | EUR | *PDGFD* | 6 | c.883C>T | p.(Arg295Cys) | D-Mis | 4.00E-06 | 35.0 | 0.56 |
| 23-025 | F | 41 | IPAH | EUR | *PDGFD* | 6 | c.926C>G | p.Ser309Cys | D-Mis | --- | 28.4 | 0.22 |
| E000820 | F | 73 | IPAH | EUR | *PDGFD* | 6 | c.961T>A | p.(Tyr321Asn) | D-Mis | 1.21E-05 | 33.0 | 0.34 |
| E010173 | F | 74 | IPAH | EUR | *PDGFD* | 6 | c.961T>A | p.(Tyr321Asn) | D-Mis | 1.21E-05 | 33.0 | 0.34 |

\*Rare, deleterious variants defined as gnomAD\_exome\_ALL AF $\leq 1.00\text{E-}04$ and LGD or missense with variable REVEL cut-off (*FBLN2* 0.92 and *PDGFD* 0.22).
\*\* Cases are heterozygous for the indicated variants.
\*\*Transcripts: *FBLN2* NM\_001998.3 and *PDGFD* NM\_033135.4

**Figure 2.**
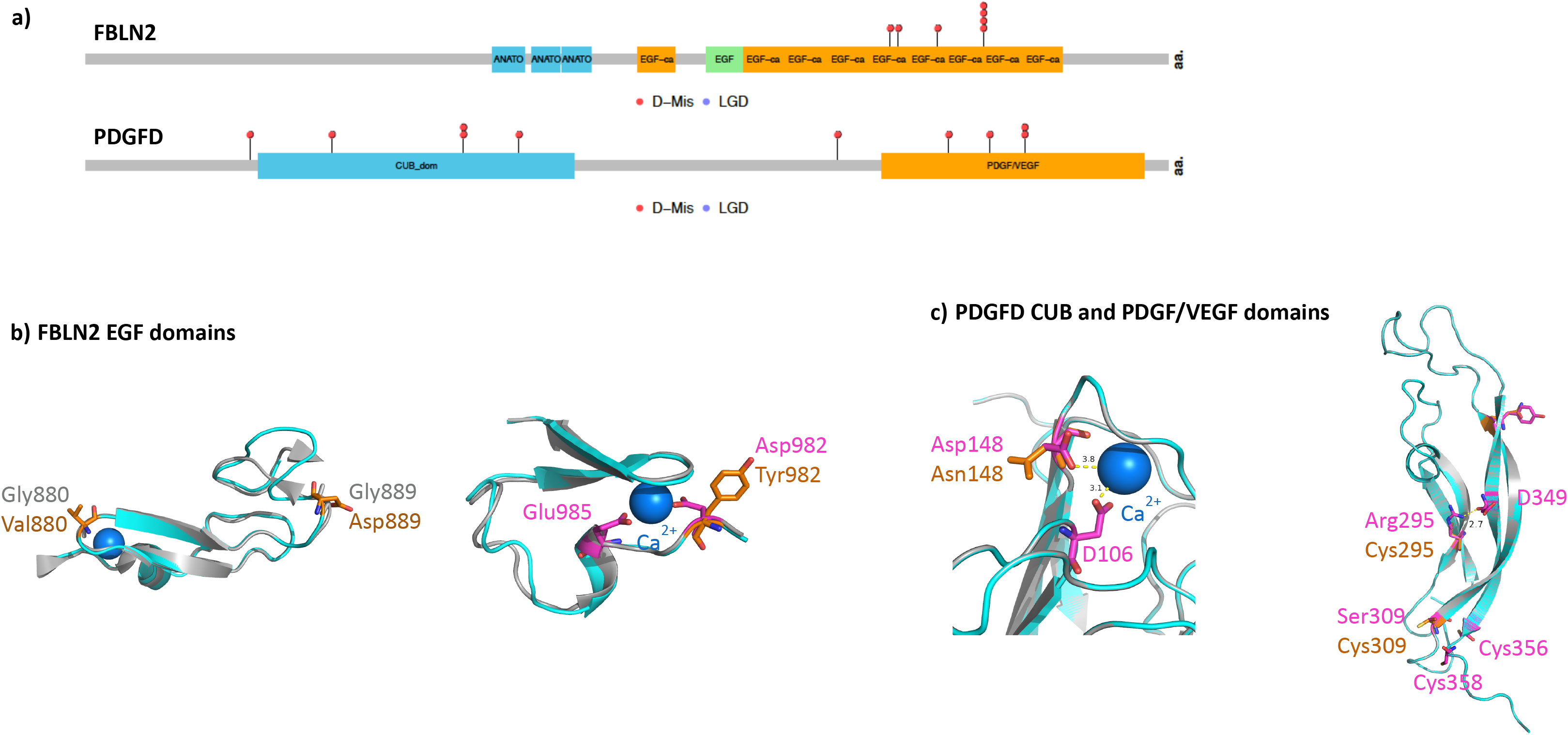
Locations of PAH-associated rare variants within FBLN2 and PDGFD protein structures. **a)** Variants and conserved domains within two-dimensional protein structures. The numbers of variants at each amino acid position is indicated along the y-axes. D-MIS, predicted deleterious missense; LGD, likely gene-disrupting (stopgain, frameshift, splicing). FBLN2: ANATO, anaphylatoxin-like 2; EGF-ca, calcium binding-endothelial growth factor-like 1; EGF, non-calcium-binding EGF domain. PDGFD: CUB, complement subcomponent; PDGF/VEGF, platelet-derived growth factor/vascular endothelial-derived growth factor domain. **b)** FBLN2 residues 858-900: p.(Gly880Val) and p.(Gly889Asp) change the conserved i+2 glycine residues of type II reverse turns within an EGF domain. Residues 981-1011: recurrent p.(Asp982Tyr) changes a residue within the highly conserved DXXE motif/calcium-binding site within an EGF domain. **c)** PDGFD residues 43-180: p.(Asp148Asn) predicted to destroy the Ca++ binding site of the CUB domain. Residues 264-364: p.(Arg295Cys) disrupts a hydrogen bond and p.(Ser309Cys) may create a new disulfide bond in the PDGF/VEGF domain.

### Clinical phenotypes of *FBLN2* and *PDGFD* variant carriers

The clinical phenotypes of all *FBLN2* and *PDGFD* variant carriers are provided in Table 3. *FBLN2* variant carriers have a similar female:male ratio (2.5:1) compared to the overall cohort (3.1:1) or IPAH alone (2.9:1). *PDGFD* variant carriers are primarily female (9:1) but the significance of this finding requires confirmation in a larger sample size. All of the *FBLN2* and *PDGFD* variant carriers have adult-onset disease, with the exception of one pediatric *PDGFD* variant carrier, with no statistically-significant differences in mean age-of-onset (53 ± 11 and 45 ± 20 years, respectively) compared to that of the overall cohort (46 ± 20 years) or IPAH alone (47 ± 20 years), excluding *FBLN2* and *PDGFD* variant carriers. *FBLN2* variant carriers exhibit a trend toward increased mean pulmonary artery pressure (62 ± 17, mmHg) and significantly increased mean pulmonary capillary wedge pressure (13 ± 2 mm Hg) compared to the overall cohort (51 ± 14, non-significant and 10 ± 4 mmHg, p = 0.15 respectively) or IPAH alone (53 ± 17, non-significant and 10 ± 24 mmHg, p = 0.01, respectively). *PDGFD* variant carriers have similar pulmonary pressures compared to the overall cohort or IPAH alone. All of the *FBLN2* and *PDGFD* variant carriers were diagnosed with WHO PAH class II or III disease and have no history of lung transplantation. Most of the *FBLN2* and *PDGFD* variant carriers have co-morbidities typical of adult IPAH patients (53, 54), including hypertension, hypothyroidism, other pulmonary diseases and metabolic diseases. Five out of seven *FBLN2* carriers have a diagnosis of systemic hypertension.

**Table 3.**
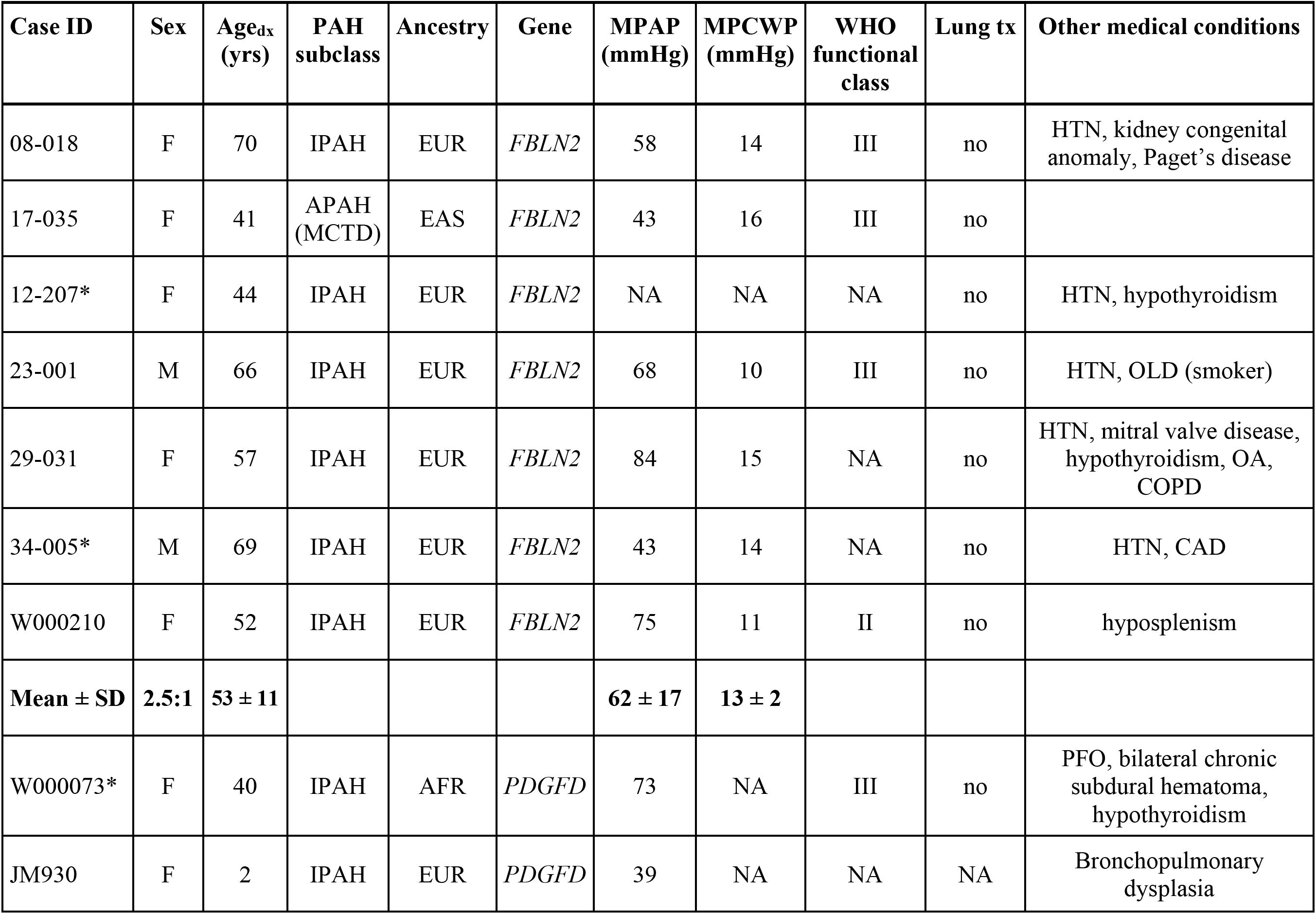
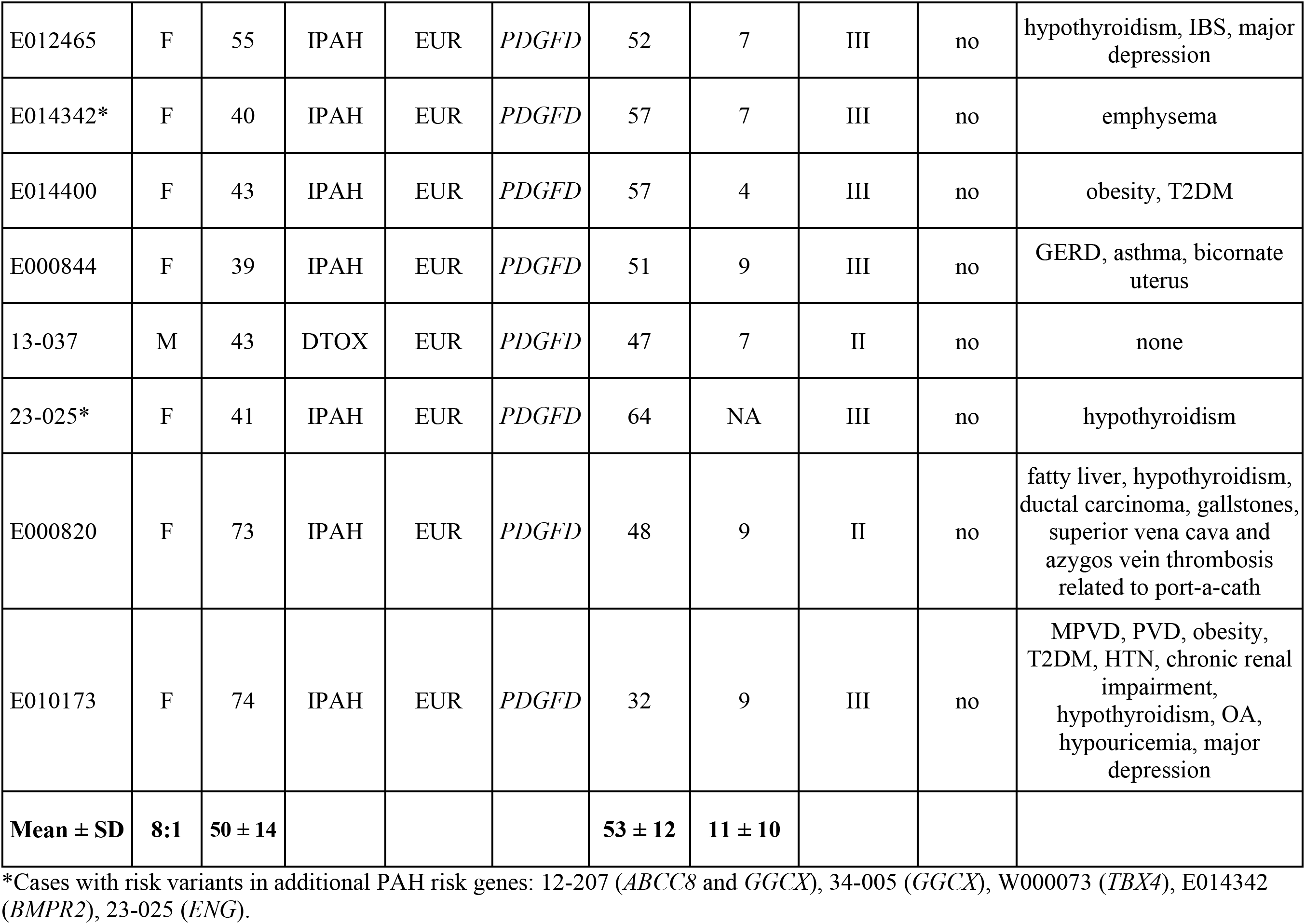
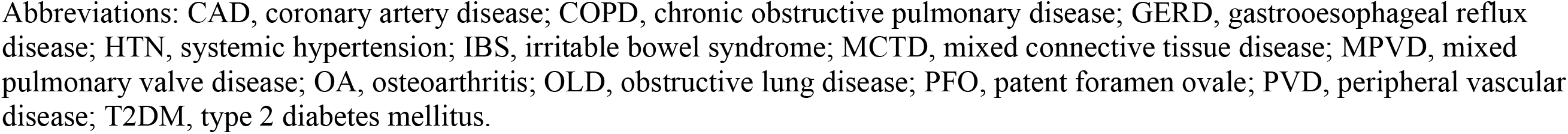
Clinical phenotypes of *FBLN2* and *PDGFD* variant carriers.

### Gene expression patterns of PAH candidate risk genes

We hypothesized that PAH risk genes are highly expressed in certain cell types relevant to the disease etiology, and that joint analysis of cell type specific expression data with genetic data could inform cell types associated with disease risk (55). We obtained single cell RNA-seq data of aorta, lung, and heart tissues available through the *Tabula Muris* project, a transcriptome compendium containing RNA-seq data from adult-staged mouse organs (48). We chose 14 tissue/cell types including endothelial, cardiac muscle, and stromal cells as a proxy for the cell types of the pulmonary artery (unavailable). A list of the tissues, cell types and the number of cells sequenced per tissue/cell type is provided in Supplementary Figure 5a. We queried gene expression for twelve known PAH risk genes (*ACVRL1*, *BMPR2*, *CAV1*, *EIF2AK4*, *ENG*, *KCNK3*, *KDR*, *NOTCH1*, *SMAD4*, *SMAD9*, *SOX17*, *TBX4*) and the two new candidate risk genes (*FBLN2*, *PDGFD*). A heat map with hierarchical clustering of relative gene expression is shown in Figure 3a. The majority of known risk genes (7/12) have relatively high expression in endothelial cells from the three tissues; most others have high expression in tissue-specific cardiac muscle, stromal cells or fibroblasts. *PDGFD* is located in the same cluster as *BMPR2*, *SOX17* and *KDR*; these genes are specifically and highly expressed in endothelial cell types. *FBLN2* is highly expressed in both endothelial and fibroblast cell types. We then randomly selected a set of 100 genes without reported associations with PAH and performed PCA of cell type specific expression profiles of known risk genes and random genes. The second component (PC2) largely separates known risk genes and random genes (Figure 3b and 3c). Consistent with hierarchical clustering, endothelial expression in all three tissues was positively correlated with PC2 (Supplementary Figure 5b). Projecting all protein-coding genes onto PC2, seven of twelve known risk genes are ranked in top 5% among all genes (Figure 3d) (binomial test: enrichment =20, p =1.6E-05). Two new candidate genes, *FBLN2* and *PDGFD*, are ranked in the top 1.8% of PC2.

**Figure 3.**
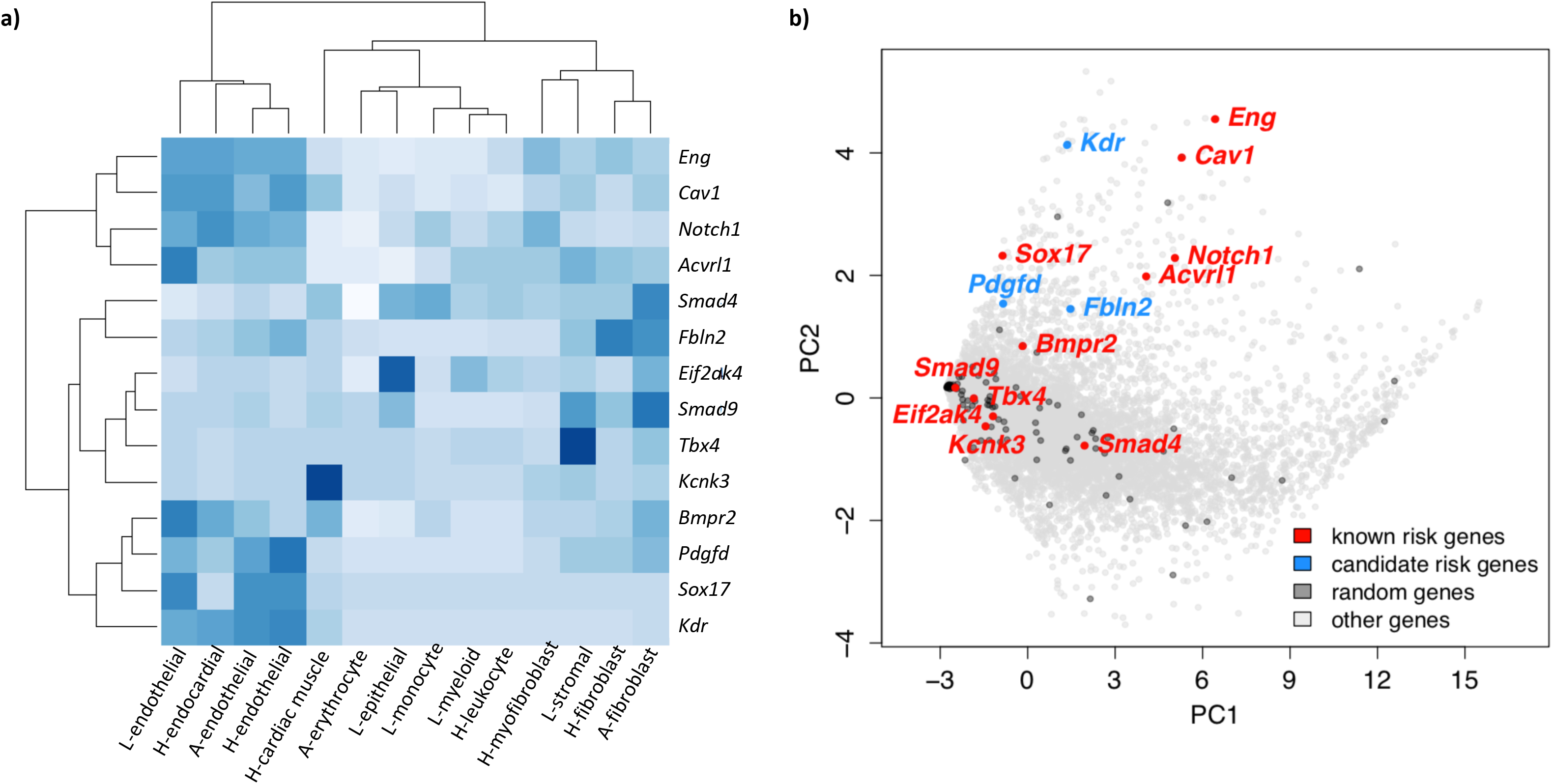
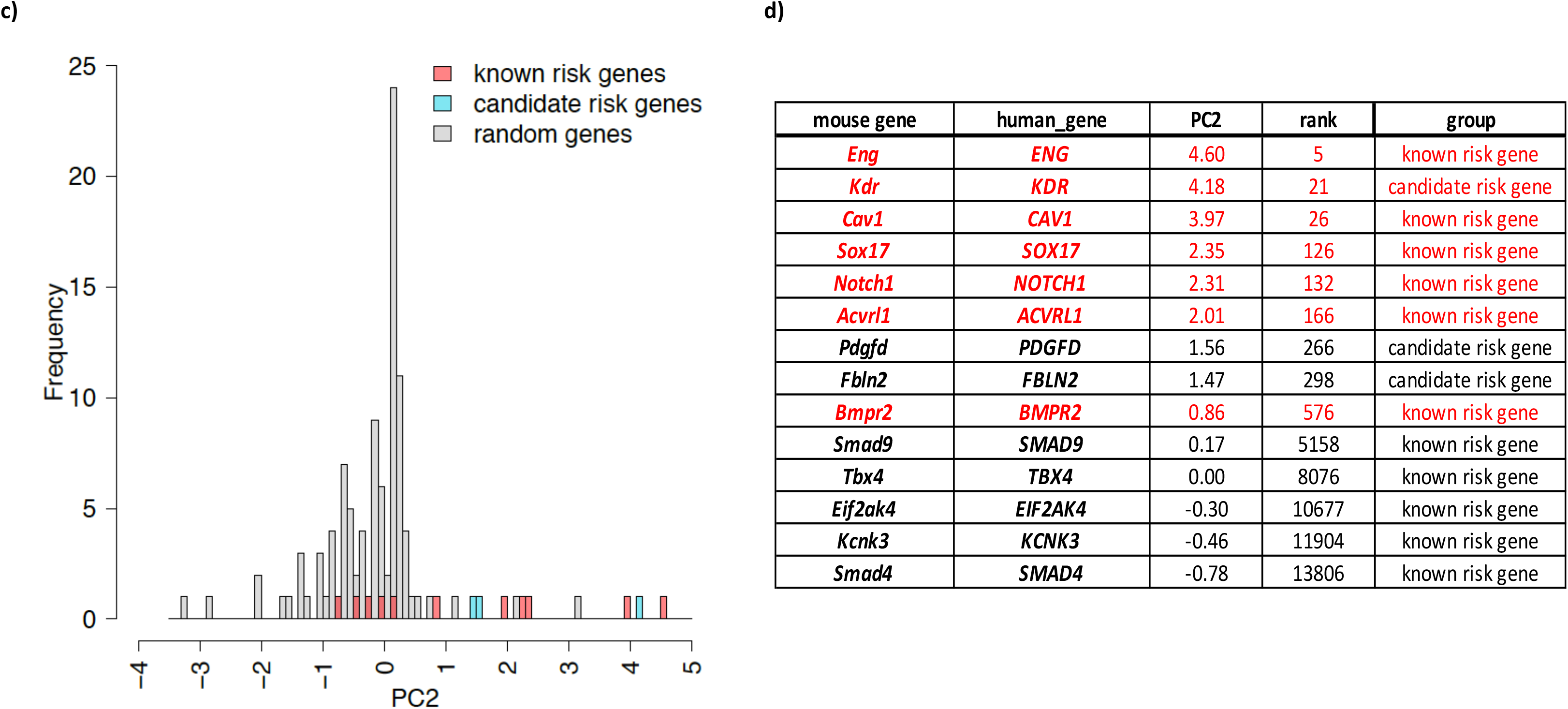
Gene expression patterns of PAH risk genes using murine single cell RNA-seq data. **a)** Heat map showing fraction of cells with >0 reads in specific cell types of lung, heart and aorta for 11 known PAH risk genes and 3 new candidate risk genes (*KDR*, *FBLN2* and *PDGFD*). L, lung; H, heart; A, aorta. **b)** PCA analysis of gene expression for PAH risk genes and a set of 100 randomly-selected genes, overlaid on a plot of all other 16,744 sequenced genes expressed in both human and mouse cells. **c)** Histogram of PC2 values for PAH risk genes and a set of 100 randomly-selected genes indicates a right shift for PC2 among PAH risk genes. **d)** Relative rank of PC2 values for PAH risk genes among 16,744 sequenced genes expressed in both human and mouse cells.

### Identification of novel candidate pediatric PAH risk genes by *de novo* variant analysis

We next focused on pediatric-onset disease, a sub-population in which genetic factors likely play a larger causal role compared to adults. The study was underpowered to carry out a gene-based case-control association analysis due to the relatively small number of pediatric patients (n = 442); however, 124 pediatric-onset PAH probands with child-parent trio data were available for *de novo* variant analysis. The trio cohort consisted mostly of IPAH (55.6%, n=66) and APAH-CHD (37.9%, n=45) cases. We performed a burden test for enrichment of *de novo* variants among all trio probands by comparing the number of variants observed vs expected based on the background mutation rate. Similar rates of *de novo* mutations were observed for synonymous, LGD alone and total missense variants (Table 4). However, there was a significant burden of D-Mis and LGD+D-Mis variants among cases over that expected by chance (Table 4). Inclusion of all protein-coding genes (n=18,939) in the burdent test identified 46 rare variants, including 30 D-Mis and 16 LGD, in cases. Confining the test to a set of 5,756 genes highly expressed in developing lung (murine E16.5 lung stromal cells) (56) or heart (murine E14.5 heart) (31), revealed a 2.5-fold enrichment of *de novo* variants among cases (n=19 D-Mis, n=30 LGD+D-Mis) over that expected by chance (p=2.0e-4, p=7.0e-6, respectively). We estimate that 18 of the variants are likely to be implicated in pediatric PAH based upon the enrichment over controls or expected by chance. Among the variants, seven are in known PAH risk genes: four in *TBX4*, two in *BMPR2*, and one in *ACVRL1*. Excluding these known risk genes, there are 23 LGD+D-Mis variants in genes highly expressed in developing heart and lung, still significantly more than expected (enrichment rate=1.95, p=0.003, 10 expected risk variants). We tested the burden of *de novo* variants among IPAH cases and observed enrichment of D-Mis and LGD+D-Mis variants similar to that of the overall trio cohort (Supplementary Table 4). The study was underpowered to detect a significant burden of *de novo* variants among APAH-CHD cases. The estimated fraction of pediatric IPAH and the overall pediatric cohort explained by *de novo* variants is 15.2% and 14.5%, respectively. A complete list of all rare, deleterious *de novo* variants carried by pediatric PAH cases is provided in Supplementary Table 5. Similar to other early-onset severe diseases, including CHD and bronchopulmonary dysplasia, the genes identified fit a general pattern for developmental disorders – genes intolerant to loss of function variants (pLI>0.5 for 40% of the genes) and with known functions as transcription factors, RNA-binding proteins, protein kinases, and chromatin modification. Three of the genes are known CHD risk genes (*NOTCH1*, *PTPN11* and *RAF1*), and 37% of the genes are known causal genes for a variety of developmental syndromes. Case variant *PTPN11* p.(Asp61Gly) is a known causal variant for Noonan syndrome(57), and *RAF1* p.Pro261 is a hotspot for multiple gain-of-function mutations, including p.(Pro261Thr), causing Noonan syndrome(58).

**Table 4.**
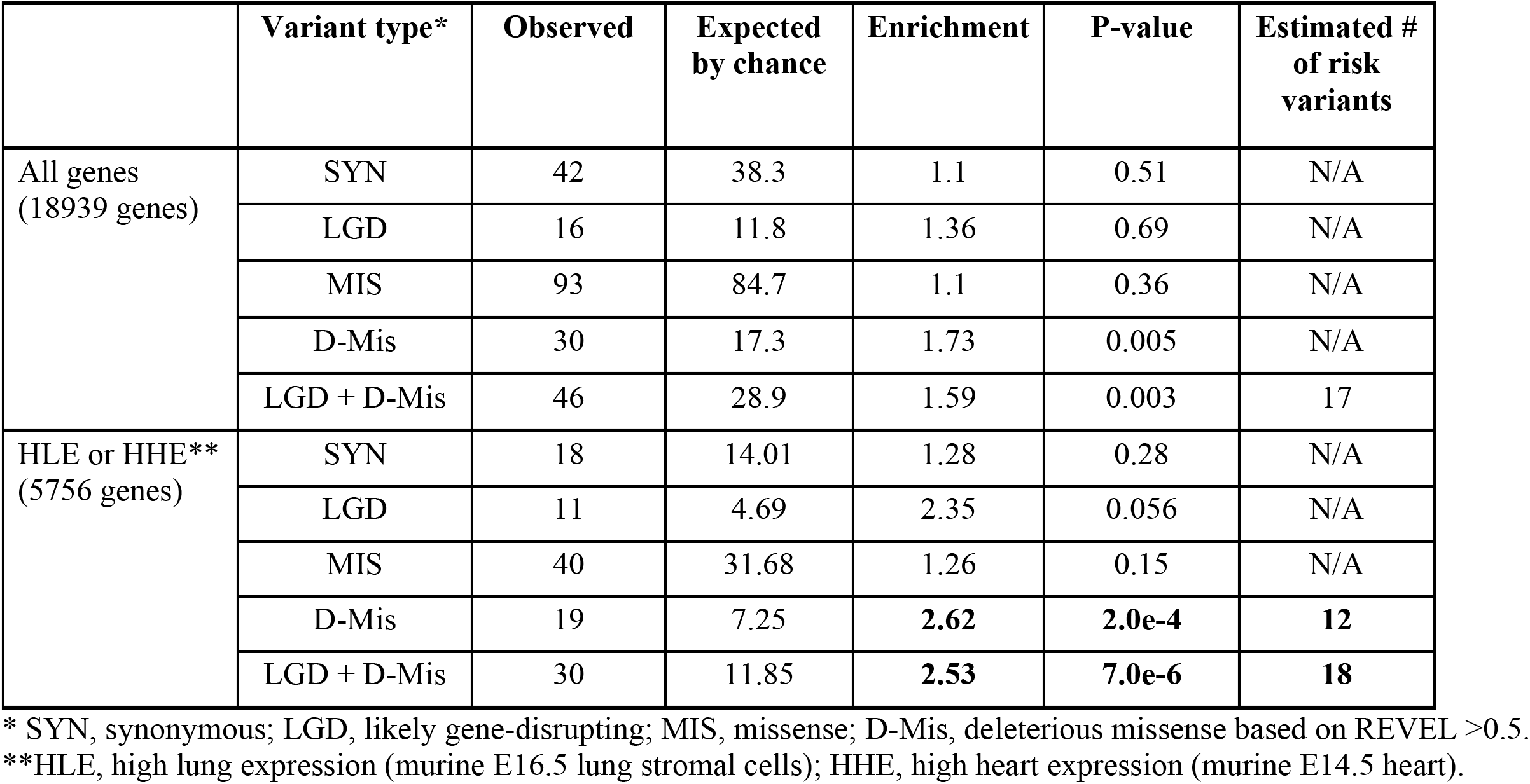
Burden of *de novo* variants in pediatric-onset PAH (n = 124 child-parent trios).

|  | Variant type* | Observed | Expected by chance | Enrichment | P-value | Estimated # of risk variants |
| --- | --- | --- | --- | --- | --- | --- |
| All genes (18939 genes) | SYN | 42 | 38.3 | 1.1 | 0.51 | N/A |
|  | LGD | 16 | 11.8 | 1.36 | 0.69 | N/A |
|  | MIS | 93 | 84.7 | 1.1 | 0.36 | N/A |
|  | D-Mis | 30 | 17.3 | 1.73 | 0.005 | N/A |
|  | LGD + D-Mis | 46 | 28.9 | 1.59 | 0.003 | 17 |
| HLE or HHE** (5756 genes) | SYN | 18 | 14.01 | 1.28 | 0.28 | N/A |
|  | LGD | 11 | 4.69 | 2.35 | 0.056 | N/A |
|  | MIS | 40 | 31.68 | 1.26 | 0.15 | N/A |
|  | D-Mis | 19 | 7.25 | **2.62** | **2.0e-4** | **12** |
|  | LGD + D-Mis | 30 | 11.85 | **2.53** | **7.0e-6** | **18** |

\* SYN, synonymous; LGD, likely gene-disrupting; MIS, missense; D-Mis, deleterious missense based on REVEL >0.5.
\*\*HLE, high lung expression (murine E16.5 lung stromal cells); HHE, high heart expression (murine E14.5 heart).

### Clinical phenotypes of pediatric *de novo* variant carriers

Among the 38 patients who carry LGD or D-Mis *de novo* variants (Supplementary Table 6), there is a 2.7:1 ratio of females to males, a mean age-of-onset of 5.6 ± 4.7 years, 54.3% of the cases (n=19) have a diagnosis of IPAH, 32.4% (n=12) APAH-CHD and an overlapping but distinct 31.4% of cases have other congenital or growth and development anomalies. *NOTCH1* variant carrier, JM1357, has a diagnosis of APAH-CHD with tetralogy of Fallot, and a recent exome sequencing study of ~800 tetralogy of Fallot cases identified *NOTCH1* as the top association signal (59). *PTPN11* variant carrier, JM155, has a diagnosis of APAH-CHD associated with Noonan syndrome and the c.182A>G variant is known to be pathogenic in Noonan syndrome. Variants in *PSMD12* cause Stankiewicz-Isidor syndrome, sometimes associated with congenital heart defects, and variant carrier 06-095 has a diagnosis of APAH-CHD. Hemodynamic data for the *de novo* variant carriers (Supplementary Table 6) was similar to that of all pediatric cases in the cohort (Supplementary Table 1).

## DISCUSSION

Combined analysis of a large US/UK cohort enriched in adult-onset IPAH cases enabled identification of five known and two new IPAH risk genes with FDR <0.1: *FBLN2* and *PDGFD* are the new genes. The association was based on a gene-level case-control analysis of 1,647 unrelated European IPAH cases. The variants contributing to *FBLN2* and *PDGFD* associations are D-Mis variants predicted to disrupt highly-conserved protein conformation, Ca^++^ binding sites or intramolecular binding sites within conserved protein domains, likely leading to important structural changes in critical domains. In addition, we confirmed the recent association of *KDR* with PAH (49, 50) based on an alternative statistical approach. We further show that all three of these candidate genes have high expression in lung and heart endothelial cell types, similar to other well established risk genes (*BMPR2* and *SOX17*), further supporting the plausibility of these genes contributing to PAH risk. *De novo* variant analysis of pediatric-onset PAH (124 trios) showed a 2.5x enrichment of rare deleterious variants, indicating that *de novo* variants contribute to ~15% of pediatric cases across PAH subtypes. The *de novo* variants implicate new candidate risk genes likely unique to pediatric PAH, but some of the molecular pathways may inform both pediatric- and adult-onset PAH.

*FBLN2* encodes an extracellular matrix protein regulating cell motility, proliferation and angiogenesis. *FBLN2* undergoes epigenetic repression or gene deletion in a variety of human cancers, indicating an important tumor suppressor role (60). Conversely, gene expression is induced during development and wound healing(61), and is required for angiotensin II-induced TGFβ signaling and cardiac fibrosis(62). Recent gene-level association of *FBLN2* with intracranial aneurysm provides additional support for a role in vascular remodeling in humans(63). We hypothesize that, in the pulmonary vasculature, gain of function variants may lead to increased TGF-β signaling, increased proliferation and medial hypertrophy. The FBLN2 protein contains 10 consecutive EGF protein-protein interaction domains, nine of which are calcium-binding. All seven of the case variants are missense variants, two of which are predicted to alter the conformation of an EGF domain and a recurrent variant carried by four cases is predicted to disrupt the Ca^++^ binding site of another EGF domain. The carriers of *FBLN2* variants have adult-onset disease with mean age-of-onset similar to the overall cohort or IPAH alone. Five of seven carriers also have a diagnosis of systemic hypertension (HTN) and it is possible that gene damaging variants in *FBLN2* contribute to the development of HTN. However, there may be other age-related genetic and environmental factors contributing to HTN. The prevalence of HTN in the overall US/UK combined cohort for adult-onset IPAH is 32%. Finally, two of our cohort cases, 08-018 and 29-031, have additional diagnoses of renal or heart anomalies, and *FLBN2* has been identified as a key regulator of development in those tissues (64–66).

*PDGFD* is a member of the *PDGF* family that function in recruiting cells of mesenchymal origin during development or to sites of injury(67). *PDGFD* is widely expressed including arterial endothelial cells, adventitial pericytes and smooth muscle cells, lung endothelial cells and smooth muscle cell progenitors of distal pulmonary arterioles. Secreted PDGFD specifically binds PDGFRβ, a widely-expressed protein that co-localizes with PDGFD in vascular smooth muscle cells. Cardiac-specific *PDGFD* transgenic mice exhibit vascular smooth muscle cell proliferation, vascular remodeling with wall thickening, severe cardiac fibrosis, heart failure and premature death(68). While effects of *Pdgfd* overexpression on the pulmonary vasculature have not been investigated, the cardiac vasculature data are consistent with a gain of function mechanism. Full length PDGFD contains two conserved protein domains, an autoinhibitory CUB domain and an enzymatic PDGF/VEGF domain; the protein undergoes proteolytic cleavage at Arg247 or Arg249 to produce an active growth factor promoting angiogenesis and vascular muscularization(69). All ten of the case variants are missense variants; four reside in the CUB domain and five reside in the active processed protein. Variant p.(Asp148Asn), carried by two patients, is predicted to disrupt the Ca^++^ binding site of the CUB domain; variants p.(Arg295Cys) and p.(Ser309Cys), carried by one and two patients respectively, are predicted to alter the conformation of the PDGF/VEGF domain. All but one of the *PDGFD* variant carriers have adult-onset disease with mean age-of-onset similar to the overall cohort or IPAH alone. Four out of ten of the *PDGFD* variant carriers have additional diagnoses of other pulmonary fibrotic and/or vascular fibrotic diseases including bronchopulmonary dysplasia, emphysema, asthma and one patient (E010173) with both mixed pulmonary valve disease and peripheral vascular disease (Table 3). Targeting the PDGF pathway with small molecule inhibitors of tyrosine kinase is an active area of investigation and several inhibitors are FDA-approved(70). Notably, imatinib reduced cardiac fibroblast proliferation and PDGFD expression 15-fold (71); data regarding effects on pulmonary arterial smooth muscle cells are warranted. A limitation of tyrosine kinase inhibitors is that they target multiple tyrosine kinases. Sequestering PDGFD with neutralizing antibodies or DNA/RNA aptamers, or preventing PDGFD-PDGFRβ interaction via oligonucleotides, may provide more specific targeting.

It is worth noting that the PAH associations of *FBLN2* and *PDGFD* were based solely on D-Mis variants. The identification of these genes was based on a method that did not rely solely on loss of function variants but also utilized accurate identification of deleterious missense variants based on implementation of a gene-specific threshold for deleteriousness predictions.

To test the plausibility of the new candidate PAH genes identified by association analysis, we leveraged publicly available single-cell RNA-seq data. *PDGFD*, and recently identified *KDR*, have very similar expression patterns as *BMPR2* and *SOX17*, two established PAH genes. PCA indicated that the PAH risk genes can largely be separated from non-risk genes based on PC2. The majority of known PAH risk genes rank in the top 5% of PC2 among 16,744 genes queried, and the new genes – *FBLN2* and *PDGFD* – rank within the top 1.8%, providing support for their candidacy as PAH risk genes. Other risk genes, like *KCNK3* and *EIF2AK4*, exert important PAH-related functions in cell types other than endothelial cells, and GDF2 is excreted from liver; thus, it will be important to consider expression patterns on a gene-specific basis. In addition, the dataset utilized in this study was based on adult-staged murine cells and is not well-suited for developmental genes such as *TBX4* and other genes likely to contribute to pediatric-onset disease. Thus, additional datasets from different time points are needed.

Five of the seventeen cases identified with rare deleterious variants in *FBLN2* or *PDGFD* also carry variants in one or two established or recently-reported risk genes. For example, participant 12-207 carries variants in *FBLN2* as well as *ABCC8* and *GGCX*, and participant W000073 carries variants in *PDGFD* and *TBX4*. Similarly, we have reported other cases from these cohorts who carry more than one rare gene variant(6, 7, 16), and eight such cases were identified in a Chinese cohort(14). This suggests that PAH cohort studies are starting to achieve adequate statistical power to not only expand the genes associated with pulmonary hypertension but also hint at the likely oligogenic nature of the disease in some individuals which is not surprising given the modest penetrance of most PAH genes. How multiple rare variants interact to affect PAH pathogenesis, penetrance or clinical outcomes will require even larger cohorts and will be one of the major aims of future large international consortia.

Our pediatric data indicate that children present with slightly higher mean pulmonary arterial pressure, decreased cardiac output and increased pulmonary vascular resistance compared to adults at diagnosis. The early age-of-onset and increased severity of clinical phenotypes suggest that there may be differences in the genetic underpinnings. *De novo* mutations have emerged as an important class of genetic factors underlying rare diseases, especially early-onset severe conditions (31, 72), due to strong negative selection decreasing reproductive fitness (73). Previously, we reported an enrichment of *de novo* variants in a cohort of 34 PAH probands with trio data (16). We have now expanded this analysis to 124 trios with pediatric-onset PAH probands and confirmed the 2.5x enrichment of *de novo* variants in cases compared to the expected rate. Seven of the variant carriers have variants in known PAH risk genes (*TBX4*, *BMPR2*, *ACVRL1*) and three of the APAH-CHD variant carriers have variants in known CHD or CHD-associated risk genes (*NOTCH1*, *PTPN11*, *PSMD12*). We previously-reported rare inherited LGD or D-Mis variants in CHD risk genes *NOTCH1* (n=5), *PTPN11* (n=1) and *RAF1* (n=2) carried by APAH-CHD cases (17). Specific inhibition of the protein encoded by *PTPN11* (SHP2) (74), and induction of mir-204 which negatively targets SHP2 (75.), improved right ventricular function in the monocrotaline rat model of PAH, suggesting a more general role of *PTPN11* in PAH.

At least eight of the other genes with case-derived *de novo* variants have plausible roles in lung/vascular development but have not been previously implicated in PAH: *AMOT* (angiomotin), *CSNK2A2* (casein kinase 2 alpha 2), *HNRNPF* (heterogeneous nuclear ribonucleoprotein F), *HSPA4* (heat shock protein family A member 4), *KDM3B* (lysine demethylase 3B), *KEAP1* (kelch-like ECH-associated protein 1), *MECOM* (MDS1 and EVI1 complex locus) and *ZMYM2* (zinc finger MIM-type containing 2). A common single nucleotide polymorphism in *MECOM* has been implicated in systemic blood pressure(76). *KEAP1* encodes the principle negative regulator of transcription factor NF-E2 p45-related factor 2 (NRF2). The NRF2-KEAP1 partnership provides an evolutionarily conserved cytoprotective mechanism against oxidative stress. Under normal conditions, KEAP1 targets NRF2 for ubiquitin-dependent degradation and represses NRF2-dependent gene expression. KEAP1 is ubiquitously expressed and aberrant oxidative stress response in the pulmonary vasculture is a recognized mechanism underlying PAH. Together, our analysis indicates that 15% of PAH cases are attributable to *de novo* variants. A larger pediatric cohort will be necessary to confirm some of these genes via replication and identify additional new genes and pathways that will likely be unique to children and not identifiable through studies of adults with PAH.

## CONCLUSIONS

We have identified *FBLN2* and *PDGFD* as new candidate risk genes for adult-onset IPAH, accounting for 0.26%, and 0.35% of 2,318 IPAH cases in the US/UK combined cohort, respectively. We note that five of seven *FBLN2* variant carriers also have a diagnosis of systemic hypertension. A few cases carry rare variants in more than one PAH risk gene, consistent with oligogenic nature of PAH in some individuals. Analysis of single-cell RNA-seq data shows that the new candidate genes have similar expression patterns to well known PAH risk genes, providing orthogonal support for the new genes and providing more mechanistic insight. We estimate that ~15% of all pediatric cases are attributable to *de novo* variants and that many of these genes are likely to have important roles in developmental processes. Larger adult and pediatric cohorts are needed to better clinically characterize these rare genetic subtypes of PAH.

## DECLARATIONS

### Ethics approval and consent to participate

This study was approved by the Institutional Review Boards (IRBs) of the Cincinnati Children’s Hospital Medical Center, the East of England Cambridge South national research ethics committee, Columbia University Irving Medical Center as well as the individual IRBs at each of the Enrolling Centers’ institutions. All patients have signed consent forms which are on file at the individual Enrolling Centers. No protected health information (PHI) on any patients enrolled in the PAH Biobank, UK NIHR Bioresource or CUIMC has been forwarded to the data analyzing group. Only the individual Enrolling Centers have the ability to re-contact any of the patients enrolled in the study. All research using these patient samples conformed to the principles of the Helsinski Declaration.

### Consent for publication

Written informed consent for publication was obtained at enrollment.

### Availability of data

The datasets used and/or analyzed during the current study are available via contact with the senior authors. For PAH Biobank data, a Confidentiality Agreement with the collaborating Regeneron Sequencing Center grants to Dr. Nichols a nonexclusive, worldwide, irrevocable, perpetual, royalty free sublicensable license to access and use the genomic data for any and all purposes. Therefore, while te PAH Biobank data are not uploaded to a publicly available database, direct access to the data are granted by the corresponding author on reasonable request who has full administrative access to all of the data. The data from the NIHRBR-RD study have been deposited in the European Genome-Phenome Archive (77). Data from most of the affected participants in the US/UK combined cohort were included in previous publications from our group (6, 7, 16, 17, 23, 50, 77). The script used for the variable threshold method is available from the following URL: https://github.com/ShenLab/VariableThresholdTest. The script used for cell type expression profiles is available from: (https://github.com/ShenLab/US_UK_combine).

### Competing interests

CG-J is a full-time employee of the Regeneron Genetics Center from Regeneron Pharmaceuticals Inc. and receives stock options as part of compensation. The remaining authors declare that they have no competing interests.

### Funding

This study was funded in part by NIH grants HL105333 (WCN, MWP), HL060056 (WKC), HL125218 (WKC, ER), and GM1200609 (YS). The UK National Cohort of Idiopathic and Heritable PAH is supported by the NIHRBR-RD, the British Heart Foundation (BHF) (SP/12/12/29836 and SP/18/10/33975), the BHF Cambridge Centre of Cardiovascular Research Excellence, and the UK Medical Research Council (MR/K020919/1), the Dinosaur Trust, BHF Programme grants to RCT (RG/08/006/25302) and NWM (RG/13/4/30107), and the UK NIHR National Institute for Health Research Cambridge Biomedical Research Centre. NWM is a BHF Professor and NIHR Senior Investigator. AL is supported by a BHF Senior Basic Science Research Fellowship (FS/13/48/30453).

All research at Great Ormond Street Hospital NHS Foundation Trust and UCL Great Ormond Street Institute of Child Health is made possible by the NIHR Great Ormond Street Hospital Biomedical Research Centre.

### Authors’ contributions

WCN, WKC, MWP, and YS had full access to all of the data in the study and take responsibility for the integrity of the data and the accuracy of the data analysis. Concept and design: WCN, WKC, SG, YS, NWM, and MWP

Acquisition, analysis, or interpretation of data: NZ, ES, MWP, CLW, JJH, XZ, YG, JK, DP, TT, KAL, ER, UK, AWC, CG-J, PAH Biobank, NIHR BioResource – Rare Diseases Cohort Study of Idiopathic and Heritable PAH, YS, WKC, NWM, SG, and WCN Drafting of the manuscript: NZ, CLW, MWP, YS, WKC, and WCN

Critical revision of the manuscript for important intellectual content: NZ, ES, MWP, CLW, JJH, XZ, YG, DP, TT, KAL, AWC, CG-J, PAH Biobank, Rare Diseases Cohort Study of Idiopathic and Heritable PAH, YS, WKC, NWM, SG, and WCN

Statistical analysis: NZ, ES, CLW, JJH, XZ, YS, SG, and WKC Supervision: YS, WKC, NWM, SG, and WCN

## Acknowledgements

Samples and/or data from the National Biological Sample and Data Repository for PAH (aka PAH Biobank) funded by an NIH investigator-initiated Resources grant (R24 HL105333 to WCN) were used in this study. We thank contributors, including the Pulmonary Hypertension Centers who collected samples used in this study, as well as patients and their families, whose help and participation made this work possible. We appreciate the contribution of the research coordinators across the clinical sites and Patricia Lanzano for coordinating the Columbia biorepository. Exome sequencing and genotyping data were generated by the Regeneron Genetics Center.

We thank NIHR BioResource volunteers for their participation, and gratefully acknowledge NIHR BioResource centers, NHS Trusts and staff for their contribution. We thank the National Institute for Health Research and NHS Blood and Transplant. The views expressed are those of the author(s) and not necessarily those of the NHS, the NIHR or the Department of Health and Social Care.

We thank the research nurses and coordinators at the specialist pulmonary hypertension centers involved in this study. We acknowledge the support of the Imperial NIHR Clinical Research Facility, the Netherlands CardioVascular Research Initiative, the Dutch Heart Foundation, Dutch Federation of University Medical Centres, the Netherlands Organization for Health Research and Development and the Royal Netherlands Academy of Sciences. We thank all the patients and their families who contributed to this research and the Pulmonary Hypertension Association (UK) for their support.

PAH Biobank Enrolling Centers’ Investigators: Russel Hirsch MD; R. James White MD, PhD; Marc Simon MD; David Badesch MD; Erika Rosenzweig MD; Charles Burger MD; Murali Chakinala MD; Thenappan Thenappan MD; Greg Elliott MD; Robert Simms MD; Harrison Farber, MD; Robert Frantz MD; Jean Elwing MD; Nicholas Hill MD; Dunbar Ivy MD; James Klinger MD; Steven Nathan MD; Ronald Oudiz MD; Ivan Robbins MD; Robert Schilz DO, PhD; Terry Fortin MD; Jeffrey Wilt MD; Delphine Yung MD; Eric Austin MD; Ferhaan Ahmad MD, PhD; Nitin Bhatt MD; Tim Lahm MD; Adaani Frost MD; Zeenat Safdar MD; Zia Rehman MD; Robert Walter MD; Fernando Torres MD; Sahil Bakshi DO; Stephen Archer MD; Rahul Argula MD; Christopher Barnett MD; Raymond Benza MD; Ankit Desai MD; Veeranna Maddipati MD.

NIHR BioResource – Rare Diseases and National Cohort Study of Idiopathic and Heritable PAH: Harm J. Bogaard, MD, PhD; Colin Church, PhD; Gerry Coghlin, MD; Robin Condliffe, MD; Mélanie Eyries, PhD; Henning Gall, MD, PhD; Stefano Ghio, MD; Barbara Girerd; PhD, Simon Holden, PhD; Luke Howard, MD, PhD; Marc Humbert, MD, PhD; David G. Kiely, MD; Gabor Kovacs, MD; Jim Lordan, PhD; Rajiv D. Machado, PhD; Robert V. MacKenzie Ross, MB, BChir; Colm McCabe, PhD; Jennifer M. Martin, MSt; Shahin Moledina, MBChB; David Montani, MD, PhD; Horst Olschewski, MD; Christopher J. Penkett, PhD; Joanna Pepke-Zaba, PhD; Laura Price, PhD; Christopher J. Rhodes, PhD; Werner Seeger, MD; Florent Soubrier, MD, PhD; Laura Southgate, PhD; Jay Suntharalingam, MD; Andrew J. Swift, PhD; Mark R. Toshner, MD; Carmen M. Treacy, BSc; Anton Vonk Noordegraaf, MD; John Wharton, PhD; Jim Wild, PhD; Stephen John Wort, PhD.

## Notes

### Competing Interest Statement

Claudia Gonzaga-Jauregui is a full-time employee of the Regeneron Genetics Center from Regeneron Pharmaceuticals Inc. and receives stock options as part of compensation. The remaining authors declare that they have no competing interests.

