## Supplemental Table 1 for "Rare variant analysis of 4,241 pulmonary arterial hypertension cases from an international consortium implicate *FBLN2*, *PDGFD* and rare *de novo* variants in PAH"

**Supplementary Table 1. Clinical characteristics and hemodynamic parameters of child- vs adult-onset PAH cases* at diagnosis.**

| Group | Age at dx (y) | F:M ratio | MPAP (mm Hg) | MCWP (mm Hg) | CO, Fisk (L/min) | PVR (Woods units) |
|---|---|---|---|---|---|---|
| Child (n=226) | 7.7 ± 5.4 (226) | 1.65:1 | 55.1 ± 18.6 (225) | 9.0 ± 3.0 (220) | 3.2 ± 1.6 (168) | 18.1 ± 11.7 (168) |
| Adult (n=2345) | 51.6 ± 14.7 (2345) | 4.02:1 | 49.6 ± 13.9 (2293) | 10.2 ± 4.2 (2231) | 4.6 ± 1.7 (1630) | 10.0 ± 5.9 (1630) |
| P-value | <0.0001** | <0.0001*** | <0.0001** | <0.0001** | <0.0001** | <0.0001** |

*Data are from the PAH Biobank (n=2,572 cases). Child-onset, <18 years of age at diagnosis.

Abbreviations: dx, diagnosis; F:M, female:male; MPAP, mean pulmonary artery pressure; MCWP, mean capillary wedge pressure; CO, cardiac output; PVR, pulmonary vascular resistance.

**Student's t-test, 2-tailed

***Fisher's exact test, 2-tailed
