## Supplemental Table 2 for "Rare variant analysis of 4,241 pulmonary arterial hypertension cases from an international consortium implicate *FBLN2*, *PDGFD* and rare *de novo* variants in PAH"

**Supplementary Table 2. Similar frequency of rare synonymous variants among European PAH cases and controls.**

| Mutation type* | PAH cases (n = 2,789) | Controls (n = 18,819) | Enrichment | P-value |
|---|---|---|---|---|
| SYN | 116,982 | 792,028 | 1.0 | 0.28 |
| MIS | 236,401 | 1,598,016 | 1.0 | 0.41 |
| Indel | 12,898 | 88,577 | 0.98 | 0.06 |

*SYN, synonymous; MIS, missense; Indel, insertion/deletion.
