## Supplemental Table 3 for "Rare variant analysis of 4,241 pulmonary arterial hypertension cases from an international consortium implicate *FBLN2*, *PDGFD* and rare *de novo* variants in PAH"

**Supplementary Table 3. Rare predicted deleterious *KDR* missense variants* among 4,175 PAH cases**.**

| Case ID | Sex | Age$_{dx}$ | PAH subclass | Ancestry | Gene *** | Exon | Nucleotide change | Amino acid change | Variant type | MAF (gnomAD exomes) | CADD score | Revel |
|---|---|---|---|---|---|---|---|---|---|---|---|---|
| 06-049 | F | 53 | APAH-CHD | EUR | *KDR* | 23 | c.3089C>G | p.(Ala1030Gly) | D-Mis | --- | 29.3 | 0.87 |
| 22-037 | F | 43 | IPAH | EUR | *KDR* | 23 | c.3175T>C | p.(Tyr1059His) | D-Mis | --- | 29.4 | 0.90 |
| E011155 | F | 74 | IPAH | EUR | *KDR* | 25 | c.3311C>A | p.(Ser1104Tyr) | D-Mis | 3.61E-05 | 29.5 | 0.86 |
| 15-032 | M | 25 | IPAH | EUR | *KDR* | 26 | c.3439C>T | p.(Pro1147Ser) | D-Mis | 9.16E-05 | 25.2 | 0.88 |
| E013241 | F | 73 | IPAH | EUR | *KDR* | 26 | c.3439C>T | p.(Pro1147Ser) | D-Mis | 9.16E-05 | 25.2 | 0.88 |
| W000314 | M | 75 | IPAH | EUR | *KDR* | 26 | c.3439C>T | p.(Pro1147Ser) | D-Mis | 9.16E-05 | 25.2 | 0.88 |

*Rare, deleterious variants defined as gnomAD_exome_ALL AF ≤1.00E-04 and LGD or missense with variable REVEL cut-off (*KDR* 0.86).

**Cases are heterozygous for the indicated variants.

***Transcript: *KDR* NM_002253.3
