## Supplemental Table 4 for "Rare variant analysis of 4,241 pulmonary arterial hypertension cases from an international consortium implicate *FBLN2*, *PDGFD* and rare *de novo* variants in PAH"

**Supplementary Table 4. Burden of *de novo* variants in pediatric-onset IPAH (n = 66 child-parent trios).**

| | **Variant type*** | **Observed** | **Expected by chance** | **Enrichment** | **P-value** | **Estimated # of risk variants** |
|---|---|---|---|---|---|---|
| All genes (18939 genes) | SYN | 18 | 20.4 | 0.88 | 0.74 | N/A |
| | LGD | 11 | 6.3 | 1.75 | 0.28 | N/A |
| | MIS | 53 | 45.1 | 1.18 | 0.23 | N/A |
| | D-Mis | 17 | 9.2 | 1.84 | 0.019 | N/A |
| | LGD + D-Mis | 28 | 15.4 | 1.82 | 0.003 | 10 |
| HLE or HHE** (5756 genes) | SYN | 10 | 7.5 | 1.34 | 0.35 | N/A |
| | LGD | 7 | 2.5 | **2.81** | **0.04** | N/A |
| | MIS | 23 | 16.9 | 1.36 | 0.14 | N/A |
| | D-Mis | 11 | 3.9 | **2.85** | **0.002** | **6** |
| | LGD + D-Mis | 18 | 6.3 | **2.86** | **0.0001** | **10** |

* SYN, synonymous; LGD, likely gene-disrupting; MIS, missense; D-Mis, deleterious missense based on REVEL >0.5.

**HLE, high lung expression (murine E16.5 lung stromal cells); HHE, high heart expression (murine E14.5 heart).
