## Supplemental Table 5 for "Rare variant analysis of 4,241 pulmonary arterial hypertension cases from an international consortium implicate *FBLN2*, *PDGFD* and rare *de novo* variants in PAH"

**Supplementary Table 5**. **Rare *de novo* LGD or D-Mis risk variants identified in 124 pediatric-onset PAH trios.**

| Gene symbol | Gene name | Transcript | Variant type | Nucleotide change | Protein change | REVEL score | AF gnomAD exomes | pLI | Lung expression (% rank) | Heart expression (% rank) | Gene-level associated medical condition(s) (OMIM #, mode of inheritance) |
|---|---|---|---|---|---|---|---|---|---|---|---|
| *ACVRL1* | Activin A receptor, type II like 1 | NM_001077401.2 | D-Mis | c.955G>C | p.(Gly319Arg) | 0.83 | . | 0.01 | 88.70 | 51.42 | Hereditary hemorrhagic telangiectasia (600376, AD) |
| *ALDH9A1* | Aldehyde dehydrogenase family 9 subfamily A member 1 | NM_000696.3 | D-Mis | c.545A>G | p.(Tyr182Cys) | 0.88 | 2.44E-05 | 0.00 | 57.11 | 59.25 | |
| *AMOT* | Angiomotin | NM_001113490.1 | LGD | c.957delC | p.(.Leu320Cysfs*55) | . | . | 0.21 | 67.96 | 94.51 | |
| *ATP6V0A2* | ATPase, H+ transporting, lysosomal, VO subunit A2 | NM_012463.4 | D-Mis | c.1184A>G | p.(Asn395Ser) | 0.82 | . | 0.00 | 70.43 | 61.38 | |
| *BMPR2* | Bone morphogenetic protein receptor, type II | NM_001204.7 | D-Mis | c.1471C>T | p.(Arg491Trp) | . | . | 1.00 | 86.44 | 42.01 | PAH (178600, AD)<br>PVOD (265450, AD) |
| *BMPR2* | Bone morphogenetic protein receptor, type II | NM_001204.7 | LGD | c.418+1G>A | p.(=) | 0.96 | . | 1.00 | 86.44 | 42.01 | PAH (178600, AD)<br>PVOD (265450, AD) |
| *BRWD3* | Bromodomain- and WD repeat-containing protein 3 | NM_153252.5 | D-Mis | c.1087G>T | p.(Asp363Tyr) | 0.58 | . | 1.00 | 61.56 | 47.34 | Mental retardation (300659, XLR) |
| *CHRNA4* | Cholinergic receptor, neuronal nicotinic, alpha polypeptide 4 | NM_000744.7 | D-Mis | c.721C>T | p.(Arg241Trp) | 0.92 | . | 0.02 | 29.90 | 18.42 | Epilepsy (600513, AD) |
| *CNTN4* | Contactin 4 | NM_001206956.1 | D-Mis | c.722A>G | p.(His241Arg) | 0.66 | . | 1.00 | 42.65 | 23.58 | |
| *CSNK2A2* | Casein kinase II, alpha 2 | NM_001896.4 | D-Mis | c.551A>T | p.(His184Leu) | 0.50 | . | 1.00 | 54.88 | 77.13 | |
| *DNMT3A* | DNA methyltransferase 3A | NM_153759.3 | D-Mis | c.473T>C | p.(Leu158Pro) | 0.78 | . | 0.00 | 98.27 | 96.73 | Acute myeloid leukemia (601626)<br>Heyn-Sproul_Jackson syndrome (618724, AD),<br>Tatton-Brown-Rahman syndrome (615879, AD) |
| *EMC8* | Endoplasmic reticulum membrane protein complex subunit 8 | NM_006067.5 | LGD | c.633G>C | p.(*211Tyrext*15) | . | 4.06E-06 | 0.26 | 72.18 | 66.27 | |
| *EMID1* | Emi domain-containing protein 1 | NM_001267895.2 | D-Mis | c.1114G>A | p.(Gly372Arg) | 0.77 | . | 0.04 | 55.13 | 36.41 | |

| Gene symbol | Gene name | Transcript | Variant type | Nucleotide change | Protein change | REVEL score | AF gnomAD exomes | pLI | Lung expression (% rank) | Heart expression (% rank) | Gene-level associated medical condition(s) (OMIM #, mode of inheritance) |
|---|---|---|---|---|---|---|---|---|---|---|---|
| *GAMT* | Guanidinoacetate methyltransferase | NM_000156.6 | D-Mis | c.490G>T | p.(Gly164Cys) | 0.92 | . | 0.00 | 42.58 | 65.43 | Cerebral creatinine deficiency syndrome 2 (612736, AR) |
| *GDPD4* | Glycerophosphodiester phosphodiesterase domain containing 4 | NM_182833.3 | LGD | c.1561T>A | p.(*521Lysext*64) | . | . | 0.00 | 9.53 | 0.00 | |
| *GRHL2* | Grainyhead-like transcription factor 2 | NM_001330593.2 | D-Mis | c.749C>T | p.(Pro250Leu) | 0.62 | . | 1.00 | 35.55 | 28.19 | Corneal dystrophy (618031, AD)<br>Deafness (608641, AD)<br>Ectodermal dysplasia/short stature syndrome (616029, AR) |
| *HNRNPF* | Heterogeneous nuclear ribonucleoprotein F | NM_001098208.1 | LGD | c.629delA | p.(Tyr210Leufs*14) | . | . | 0.86 | 85.44 | 97.61 | |
| *HSPA4* | Heat shock protein family A (HSP70) member 4 | NM_002154.4 | D-Mis | c.2051C>G | p.(Pro684Arg) | 0.62 | 4.10E-06 | 0.03 | 42.59 | 95.61 | |
| *ITPR1* | Inositol 1,4,5-triphosphate receptor type 1 | NM_001168272.1 | D-Mis | c.3614C>T | p.(Ala1205Val) | 0.69 | 1.69E-05 | 1.00 | 94.57 | 78.37 | Gillespie syndrome (206700, AD, AR)<br>Spinocerebellar ataxia (606658, AD; 117360, AD) |
| *KDM3B* | Lysine demethylase 3B | NM_016604.4 | D-Mis | c.3298C>T | p.(Pro1100Ser) | 0.66 | . | 1.00 | 89.37 | 86.59 | |
| *KEAP1* | Kelch-like ECH-associated protein 1 | NM_012289.4 | LGD | c.1752C>A | p.(Tyr584*) | . | . | 0.25 | 79.29 | 82.09 | |
| *MANEA* | Mannosidase, endo-alpha | NM_024641.4 | LGD | c.678_679insTTCTTGG | p.(Ser227Phefs*7) | . | . | 0.00 | 32.78 | 54.66 | |
| *MBTPS1* | Membrane-bound transcription factor protease, site 1 | NM_003791.4 | D-Mis | c.1342G>A | p.(Ala448Thr) | 0.61 | 5.41E-06 | 0.13 | 97.85 | 95.59 | Spondyloepiphyseal dysplasia (618392, AR, 1 patient only) |
| *MECOM* | MDS1 and EVI1 complex locus | NM_001163999.1 | D-Mis | c.2285T>C | p.(Phe762Ser) | 0.76 | . | 1.00 | 81.79 | 59.65 | Radioulnar synostosis and amegakaryocytic thrombopenia (RUSAT2, 616738, AD) |
| *MFN2* | Mitofusin 2 | NM_001127660.1 | D-Mis | c.311G>A | p.(Arg104Gln) | 0.94 | . | 1.00 | 76.46 | 96.94 | Charcot-Marie-Tooth disease (609260, AD; 617087, AR)<br>Hereditary motor and sensory neuropathy (601152, AD) |
| *MYOM1* | Myomesin 1 | NM_019856.2 | LGD | c.3019C>T | p.(Arg1007*) | . | 1.27E-05 | 0.00 | 57.08 | 99.42 | |

| Gene symbol | Gene name | Transcript | Variant type | Nucleotide change | Protein change | REVEL score | AF gnomAD exomes | pLI | Lung expression (% rank) | Heart expression (% rank) | Gene-level associated medical condition(s) (OMIM #, mode of inheritance) |
|---|---|---|---|---|---|---|---|---|---|---|---|
| *NOTCH1* | Notch receptor 1 | NM_017617.5 | D-Mis | c.1430T>A | p.(Ile477Asn) | 0.74 | . | 1.00 | 87.73 | 87.89 | Adams-Oliver syndrome (616028AD)<br>Aortic valve disease (109730, AD) |
| *NUCB1* | Nucleobindin 1 | NM_006184.6 | LGD | c.568dupT | p.(Tyr190Leufs*29) | . | . | 0.00 | 94.51 | 93.26 | |
| *OLFML2B* | Olfactomedin-like 2B | NM_001297713.1 | D-Mis | c.1694T>A | p.(Ile565Asn) | 0.69 | . | 0.00 | 74.96 | 75.85 | |
| *PSMD12* | Proeasome 26S subunit, non-ATPase 12 | NM_174871.4 | LGD | c.1207_1209del | p.(Asn403del) | . | . | 1.00 | 84.88 | 89.07 | Stankiewicz-Isidor syndrome (617516, AD) |
| *PTPN11* | Protein tyrosine phosphatase non-receptor type 11 | NM_002834.5 | D-Mis | c.182A>G | p.(Asp61Gly) | 0.92 | . | 1.00 | 92.32 | 94.20 | LEAPARD syndrome (151100, AD)<br>Leukemia (607785)<br>Metachondromatosis (156250, AD)<br>Noonan syndrome (163950, AD) |
| *PTPRK* | Protein tyrosine phosphatase receptor type K | NM_001291984.2 | D-Mis | c.4202T>C | p.(Val1401Ala) | 0.54 | . | 0.98 | 93.06 | 90.50 | |
| *RAF1* | Raf-1 proto-oncogene, serine/threonine kinase | NM_002880.3 | D-Mis | c.781C>A | p.(Pro261Thr) | 0.87 | . | 1.00 | 26.87 | 91.43 | Cardiomyopathy (615916, AD)<br>LEOPARD syndrome (611554)<br>Noonan syndrome (611553, AD) |
| *RALGAPA1* | Ral GTPase-activating protein catalytic subunit A1 | NM_001283043.3 | LGD | c.2022_2023insCAATTATTTAAAC | p.(Arg675Glnfs*9) | . | . | 1.00 | 90.19 | 73.73 | Neurodevelopmental disorder with neonatal espiratory insufficiency and thermodysregualtion (618797, AR) |
| *RASA2* | RAS P21 protein activator 2 | NM_001303245.2 | D-Mis | c.1916C>T | p.(Thr639Ile) | 0.56 | 1.63E-05 | 0.00 | 74.96 | 66.34 | |
| *SLC25A24* | Solute carrier family 25 member 24 | NM_013386.5 | D-Mis | c.649C>T | p.(Arg217Cys) | 0.81 | . | 0.00 | 77.16 | 73.97 | Fontaine progeroid syndrome (612289, AD) |
| *SLC38A6* | Solute carrier family 38 member 6 | NM_153811 | LGD | c.364-1G>T | p.(=) | . | . | 0.00 | 44.38 | 33.57 | |
| *SRPRA* | SPR receptor subunit alpha | NM_001177842.1 | D-Mis | c.100C>A | p.(Arg34Ser) | 0.64 | . | NA | NA | NA | |
| *TBX4* | T-box transcription factor 4 | NM_018488.3 | LGD | c.293C>G | p.(Pro98Arg) | 0.97 | . | 0.41 | 98.98 | 24.64 | Ischiocoxopodopatellar syndrome with or without PAH (147891, AD) |

| Gene symbol | Gene name | Transcript | Variant type | Nucleotide change | Protein change | REVEL score | AF gnomAD exomes | pLI | Lung expression (% rank) | Heart expression (% rank) | Gene-level associated medical condition(s) (OMIM #, mode of inheritance) |
|---|---|---|---|---|---|---|---|---|---|---|---|
| | | | | | | | | | | | Posterior amelia with pelvic and pulmonary hypoplasia syndrome (601360, AR) |
| *TBX4* | T-box transcription factor 4 | NM_018488.3 | LGD | c.538_547del | p.(Pro180Ilefs*45) | . | . | 0.41 | 98.98 | 24.64 | Ischiocoxopodopatellar syndrome with or without PAH (147891, AD) Posterior amelia with pelvic and pulmonary hypoplasia syndrome (601360, AR) |
| *TBX4* | T-box transcription factor 4 | NM_018488.3 | D-Mis | c.985G>T | p.(Asp329Tyr) | 0.61 | . | 0.41 | 98.98 | 24.64 | Ischiocoxopodopatellar syndrome with or without PAH (147891, AD) Posterior amelia with pelvic and pulmonary hypoplasia syndrome (601360, AR) |
| *TBX4* | T-box transcription factor 4 | NM_018488.3 | LGD | c.1054C>T | p.(Arg352*) | . | . | 0.41 | 98.98 | 24.64 | Ischiocoxopodopatellar syndrome with or without PAH (147891, AD) Posterior amelia with pelvic and pulmonary hypoplasia syndrome (601360, AR)Posterior amelia with pelvic and pulmonary hypoplasia syndrome(601360) |
| *TRH* | Thyrotropin-releasing hormone | NM_007117.5 | D-Mis | c.253C>A | p.(His85Asn) | 0.54 | . | 0.00 | 12.25 | 75.66 | Thyrotropin-releasing hormone deficiency (275120, AR) |
| *TUBB6* | Tubulin beta 6 class V | NM_001303527.2 | DMis | c.40C>T | p.(Arg14Trp) | 0.68 | 8.13E-06 | 0.00 | 92.18 | 89.57 | Congenital facial palsy with ptosis and velopharyngeal dysfunction (617732, AD) |
| *ZMYM2* | Zinc finger MYM-type containing 2 | NM_001190965.3 | LGD | c.1618C>T | p.(Arg540*) | . | . | 0.97 | 93.10 | 77.22 | |
| *ZNF620* | Zinc finger protein 620 | NM_175888.4 | LGD | c.74G>A | p.(Trp25*) | . | . | 0.00 | NA | NA | |

*Rare, deleterious variants defined as gnomAD AF ≤1.00E-04 and LGD or missense with REVEL <0.5.
