## Supplemental Table 6 for "Rare variant analysis of 4,241 pulmonary arterial hypertension cases from an international consortium implicate *FBLN2*, *PDGFD* and rare *de novo* variants in PAH"

**Supplementary Table 6. Clinical characteristics of pediatric PAH cases with rare *de novo* LGD or D-Mis variants.**

| Gene symbol | Variant type | Gene-level associated medical condition(s) | Case ID | Sex | Age of dx | Genetic ancestry | PAH class | PAH subclass | Heart defect | Growth & development phenotype | Other medical condition(s) | MPAP (mmHg) | MPCWP (mmHg) | CO Fick (L/min) | PVR (Woods units) | Vital status |
|---|---|---|---|---|---|---|---|---|---|---|---|---|---|---|---|---|
| *ACVRL1* | D-Mis | HHT | JM0057 | M | 6 | AFR | APAH | HHT | | | NA | 54 | 4 | 2.7 | 18.5 | Deceased |
| *ALDH9A1, TUBB6* | D-Mis | *TUBB6*: Congenital facial palsy with ptosis and velopharyngeal dysfunction | 15-002* | F | 9 | AFR | IPAH | | | | | 75 | 8 | 5.8 | 11.55 | Transfer to adult care |
| *AMOT, ZNF620* | LGD | | JM0004* | M | 1 | EUR | IPAH | | | | | 36 | 11 | 3.6 | 6.94 | Alive 2016 |
| *ATP6V0A2* | D-Mis | | JM1344 | M | 12.5 | EAS | IPAH | | | Autism | | 54 | 9 | 2.4 | 12.08 | Alive 2020 |
| *BMPR2* | D-Mis | PAH; PVOD | JM625 | M | 3 | EUR | IPAH | | | | Anemia, depression, renal failure | NA | NA | NA | NA | NA |
| *BMPR2, MECOM* | LGD | *BMPR2*: PAH; PVOD *MECOM*: RUSAT2 | LP2001061* | F | 3 | EUR | IPAH | | | | | NA | NA | NA | NA | NA |
| *BRWD3, GDPG4* | D-Mis, LGD | *BRWD3*: Mental retardation | JM140* | F | 3 | EUR | APAH | CHD | TOGV | | Asthma, depression | 59 | 14 | 6.8 | 6.62 | Deceased (17 y) |
| *CHRNA4* | D-Mis | Epilepsy | JM187 | M | 0.75 | EUR | APAH | CHD | VSD, coarctation of the aorta | Multiple congenital anomalies | | NA | NA | NA | NA | NA |
| *CNTN4* | D-Mis | | JM0035 | F | 12 | EUR | IPAH | | | | | 29 | 5 | 3.8 | 6.32 | Deceased following tx |
| *CSNK2A2, SLC38A6, TRH* | D-Mis, LGD, D-Mis | *TRH*: TRH deficiency | JM0028* | F | 9 | EUR | IPAH | | | | | 37 | 8 | 2 | 14.5 | Deceased |

| Gene symbol | Variant type | Gene-level associated medical condition(s) | Case ID | Sex | Age of dx | Genetic ancestry | PAH class | PAH subclass | Heart defect | Growth & development phenotype | Other medical condition(s) | MPAP (mmHg) | MPCWP (mmHg) | CO Fick (L/min) | PVR (Woods units) | Vital status |
|---|---|---|---|---|---|---|---|---|---|---|---|---|---|---|---|---|
| *DNMT3A* | D-Mis | Heyn-Sproul_Jackson syndrome, Tatton-Brown-Rahman syndrome, AML | JM217 | F | 15 | EUR | APAH | CHD | VSD, Eisenmenger syndrome | Down syndrome | Asthma, thyroid cancer | NA | NA | NA | NA | NA |
| *EMC8* | LGD | | JM1307 | M | 3 | SAS | IPAH | | | NA | NA | NA | NA | NA | NA | NA |
| *EMID1* | D-Mis | | 15-054 | M | 4 | AFR | IPAH | | | | | 34 | 9 | NA | NA | NA |
| *GAMT, MFN2* | D-Mis | *GAMT*: Cerebral creatinine deficiency syndrome 2 *MFN*: Charcot-Marie-Tooth disease, Hereditary motor and sensory neuropathy | FPPH4004* | M | 10 | EUR | FPAH | | | NA | NA | NA | NA | NA | NA | NA |
| *GRHL2* | D-Mis | Corneal dystrophy, Deafness, Ectodermal dysplasia/short stature syndrome | LP2000991 | M | 2 | EUR | IPAH | | secundum ASD | | | 69 | NA | NA | NA | NA |
| *HNRNPF* | LGD | | FPPH5703 | F | 0.25 | EUR | FPAH | | | PPHN, myelodysplastic syndrome | Microangiopathic hemolytic anemia, thrombocytopenia | 56 | 3 | 1.4 | 37.86 | Deceased (1.25 y) |
| *HSPA4* | D-Mis | | JM192 | F | 15 | EUR | APAH | CHD | ASD, PDA | | | NA | 6 | 12.9 | 0.93 | NA |
| *ITPR1* | D-Mis | | 15-042 | M | 5 | EUR | IPAH | | | | | 106 | 5 | 5.6 | 18.04 | Alive 2018 |
| *KDM3B* | D-Mis | | 15-006 | F | 8 | EUR | IPAH | | | | | 37 | 5 | NA | NA | Alive 2018 |

| **Gene symbol** | **Variant type** | **Gene-level associated medical condition(s)** | **Case ID** | **Sex** | **Age of dx** | **Genetic ancestry** | **PAH class** | **PAH subclass** | **Heart defect** | **Growth & development phenotype** | **Other medical condition(s)** | **MPAP (mmHg)** | **MPCWP (mmHg)** | **CO Fick (L/min)** | **PVR (Woods units)** | **Vital status** |
|---|---|---|---|---|---|---|---|---|---|---|---|---|---|---|---|---|
| *KEAP1* | LGD | | JM852 | F | 2 | EUR | IPAH | | | Developmental delay, Incontinentia pigmenti, spastic diplegia | | 63 | 11 | 1.5 | 34.67 | Deceased |
| *MANEA, RALGAPA1* | LGD | *MANEA*: Gillespie syndrome, Spinocerebellar ataxia *RALGAPA1*: Neurodevelopmental disorder with neonatal respiratory insufficiency and thermodysregualtion | JM0010* | F | 11 | EUR | IPAH | | | | | 57 | 8 | 5.7 | 8.6 | Deceased (25 y) |
| *MBTPS1* | D-Mis | Spondyloepiphyseal dysplasia (1 patient) | JM0024 | F | 4 | EUR | APAH | CHD | ASD, dextrocardia | Small stature for age | | 52 | 9 | 2.2 | 19.55 | Alive 2020 |
| *MYOM1* | LGD | | JM1367 | F | 8 | EUR | IPAH | | Secundum ASD | Chronic lung disease of prematurity | | NA | 7 | NA | NA | Alive 2020 |
| *NOTCH1* | D-Mis | Adams-Oliver syndrome Aortic valve disease | JM1357 | F | 1 | EUR | APAH | CHD | TOF | Failure to thrive | | 52 | 8 | NA | NA | Deceased (11 y) |
| *NUCB1* | LGD | | JM171 | F | 5 | EUR | IPAH | | | NA | NA | NA | NA | NA | NA | NA |
| *OLFML2B, RAF1* | D-Mis | *RAF1*: Cardiomyopathy LEOPARD syndrome Noonan syndrome | JM1088* | F | 13 | EUR | IPAH | | | Failure to thrive | Myxomatous AV valve (neoplasm) | NA | NA | NA | NA | NA |
| *PSMD12* | LGD | Stankiewicz-Isidor syndrome | 06-095 | F | 1 | EUR | APAH | CHD | PDA | | | 65 | 5 | 2 | 30.00 | NA |

| Gene symbol | Variant type | Gene-level associated medical condition(s) | Case ID | Sex | Age of dx | Genetic ancestry | PAH class | PAH subclass | Heart defect | Growth & development phenotype | Other medical condition(s) | MPAP (mmHg) | MPCWP (mmHg) | CO Fick (L/min) | PVR (Woods units) | Vital status |
|---|---|---|---|---|---|---|---|---|---|---|---|---|---|---|---|---|
| *PTPN11* | D-Mis | LEAPARD syndrome<br>Leukemia<br>Metachondromatosis<br>Noonan syndrome | JM155 | F | 6 | EUR | APAH | CHD | ASD | Noonan syndrome | | 46 | 8 | 3.7 | 10.27 | NA |
| *PTPRK* | D-Mis | | JM200 | F | 2 | EUR | APAH | CHD | ASD, PDA, TOGV, VSD | | | 82 | 5 | 1.7 | 45.29 | Deceased (9 y) |
| *RASA2* | D-Mis | | JM138 | F | 0.42 | AMR | APAH | CHD | PDA | Congenital diplegia, small stature for age | | 51 | 8 | NA | NA | NA |
| *SLC25A24* | D-Mis | Fontaine progeroid syndrome | JM216 | F | 2 | EUR | IPAH | | | | Cystic fibrosis | 39 | 11 | 3.1 | 9.03 | Alive 2020 |
| *SRPRA* | D-Mis | | FPPH133-01 | F | 15 | EUR | FPPH | HHT | | Autism | Obsessive compulsive disorder | 69 | 10 | 3.2 | 18.44 | Deceased |
| *TBX4* | D-Mis | Ischiocoxopodopatellar syndrome with or without PAH<br>Posterior amelia with pelvic and pulmonary hypoplasia syndrome | JM0002 | F | 2 | Unknown | APAH | CHD | NA | | | | | | | |
| *TBX4* | D-MIS | Ischiocoxopodopatellar syndrome with or without PAH<br>Posterior amelia with pelvic and pulmonary hypoplasia syndrome | 01-008 | F | 7 | EUR | IPAH | | | | | 40 | 8 | 4.8 | 6.67 | |
| *TBX4* | LGD | Ischiocoxopodopatellar syndrome with or without PAH | FPPH9002 | M | 2 | EUR | FPAH | Portal | | NA | NA | NA | NA | NA | NA | NA |

| Gene symbol | Variant type | Gene-level associated medical condition(s) | Case ID | Sex | Age of dx | Genetic ancestry | PAH class | PAH subclass | Heart defect | Growth & development phenotype | Other medical condition(s) | MPAP (mmHg) | MPCWP (mmHg) | CO Fick (L/min) | PVR (Woods units) | Vital status |
|---|---|---|---|---|---|---|---|---|---|---|---|---|---|---|---|---|
| | | Posterior amelia with pelvic and pulmonary hypoplasia syndrome | | | | | | | | | | | | | | |
| *TBX4* | LGD | Ischiocoxopodopatellar syndrome with or without PAH Posterior amelia with pelvic and pulmonary hypoplasia syndrome | JM847 | M | 1-day | EUR | APAH | CHD | Alveolar hypoplasia | | | 66 | 10 | NA | NA | NA |
| *ZMYM2* | LGD | | JM630 | M | 3 | EUR | IPAH | | | Actelectasis, bilateral lung; traction bronchiectasis; rib irregularities, bilateral; idiopathic scoliosis | | 61 | 13 | 2.8 | 17.14 | Alive 2020 |

Abbreviations: AML, acute myeloid leukemia; RUSAT2, Radioulnar synostosis and amegakaryocytic thrombopenia; TRH, thyroid hormone deficiency.

*Multiple *de novo* variants identified in patient 15-002 (*ALDH9A1*, *TUBB6*), patient LP2001061 (*BMPR2* and *MECOM*), patient FPPH4004 (*GAMT*, *MFN2*), patient JM0004 (*AMOT*, *ZNF620*), patient JM0010 (*MANEA*, *RALGAPA1*), patient JM0028 (*CSNK2A2*, *TRH*), patient JM140 (*BRWB3*, *GDPD4*), patient JM1088 (*OLFML2B*, *RAF1*).
