## Supplemental Figure 1 for "Rare variant analysis of 4,241 pulmonary arterial hypertension cases from an international consortium implicate *FBLN2*, *PDGFD* and rare *de novo* variants in PAH"

**Supplementary Figure 1.** Gene-level burden test for rare synonymous variants using 2789 European PAH cases and 18,819 European controls (11,101 SPARK parents, 7718 NFE gnomAD v2.1.1). Results of a binomial test confined to rare synonymous variants in 20,000 protein-coding genes.

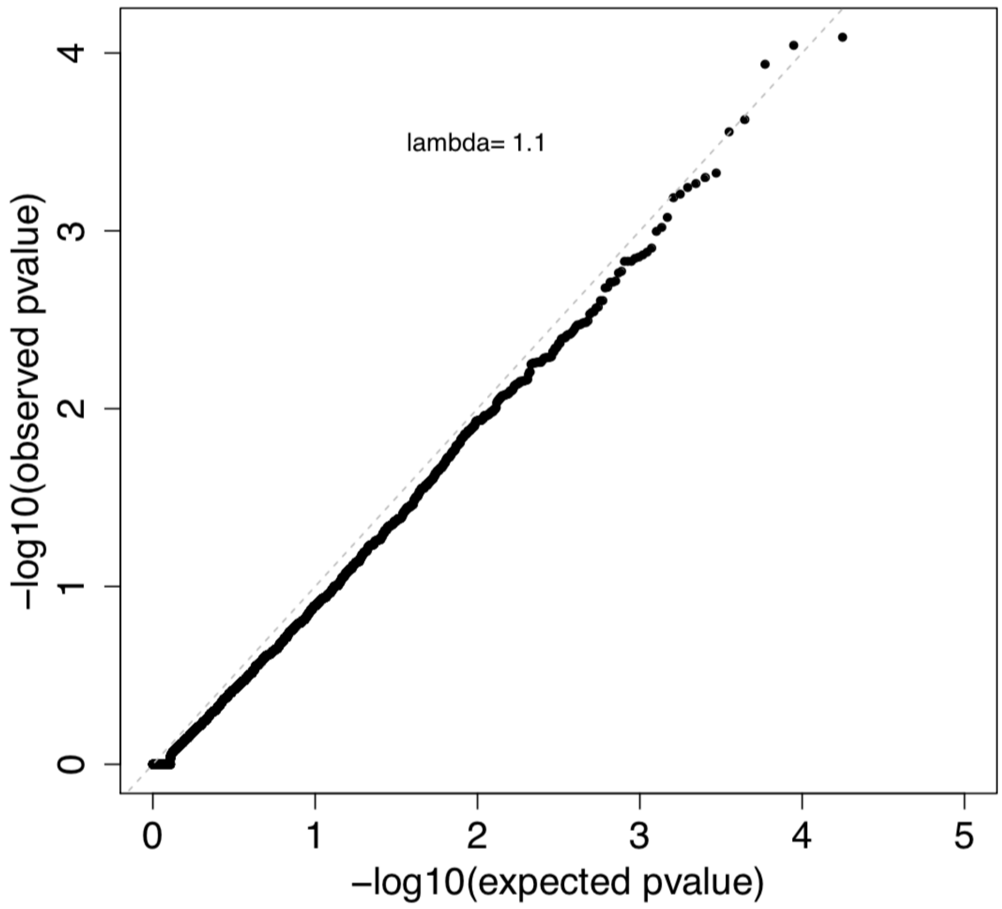
