## Supplemental Figure 2 for "Rare variant analysis of 4,241 pulmonary arterial hypertension cases from an international consortium implicate *FBLN2*, *PDGFD* and rare *de novo* variants in PAH"

**Supplementary Figure 2. Gene-based association analysis using 2,789 European cases from all PAH subclasses and 18,819 European controls (11,101 SPARK parents, 7,718 NFE gnomAD v2.1.1). a)** Results of a binomial test confined to rare, likely gene damaging (LGD) and predicted deleterious missense (D-Mis) variants or D-Mis only variants in 20,000 protein-coding genes. Horizontal gray line indicates the Bonferroni-corrected threshold for significance. **b)** Complete list of top association genes (FDR<0.1).

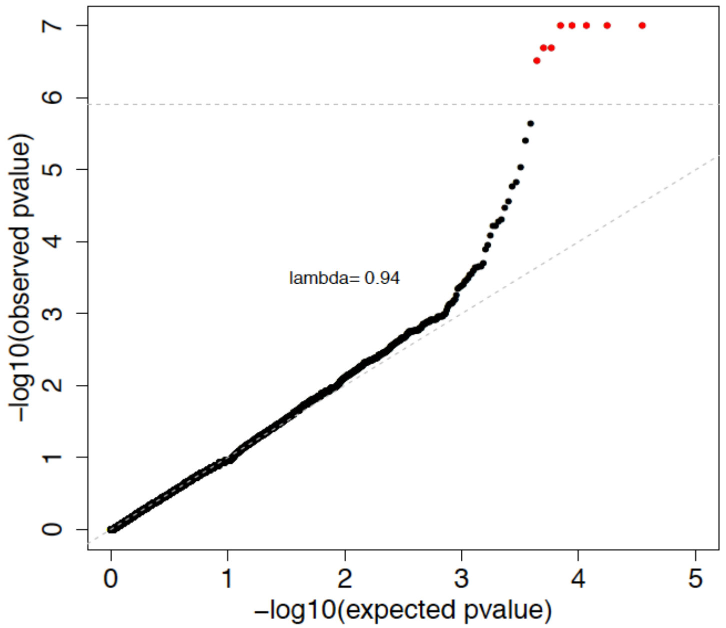


| Gene | Gene name | Revel cut-off | Variant type | P-value | FDR |
|---|---|---|---|---|---|
| *BMPR2* | Bone morphogenetic receptor type 2 | 0.68 | LGD + D-Mis | 1.00E-07 | 7.20E-04 |
| *BMPR2* | Bone morphogenetic receptor type 2 | 0.68 | D-Mis only | 1.00E-07 | 7.20E-04 |
| *TBX4* | T-box transcription factor 4 | 0.96 | LGD + D-Mis | 1.00E-07 | 7.20E-04 |
| *GDF2* | Growth differentiation factor 2 | 0.72 | LGD + D-Mis | 1.00E-07 | 7.20E-04 |
| *GDF2* | Growth differentiation factor 2 | 0.72 | D-Mis only | 1.00E-07 | 7.20E-04 |
| *KDR* | Kinase insert domain receptor | 0.86 | LGD + D-Mis | 2.00E-07 | 1.03E-03 |
| *ACVRL1* | Activin receptor like 1 | 0.66 | LGD + D-Mis | 2.00E-07 | 1.03E-03 |
| *SOX17* | SRY box transcription factor 17 | 0.80 | LGD + D-Mis | 3.00E-07 | 1.35E-03 |
| *ACVRL1* | Activin receptor like 1 | 0.66 | D-Mis only | 2.30E-06 | 0.01 |
| *COL6A5* | Collagen type VI alpha 5 chain | 0.70 | LGD + D-Mis | 4.00E-06 | 0.01 |
| *AQP1* | Aquaphorin 1 | 0.76 | LGD + D-Mis | 1.50E-05 | 0.04 |
| *ATP13A3* | ATPase type 13A3 | 1.0 | LGD + D-Mis | 1.70E-05 | 0.05 |
| *JPT2* | Jupiter microtubule-associated homolog 2 | 0.22 | D-Mis only | 2.80E-05 | 0.07 |
| *FBLN2* | Fibulin 2 | 0.92 | D-Mis only | 3.40E-05 | 0.08 |
