## Supplemental Figure 3 for "Rare variant analysis of 4,241 pulmonary arterial hypertension cases from an international consortium implicate *FBLN2*, *PDGFD* and rare *de novo* variants in PAH"

**Supplementary Figure 3. Depth of coding sequence coverage.** Percentage of case and control samples with >10X or >15X coverage for novel PAH genes **a)** *FBLN2* and **b)** *PDGFD*.

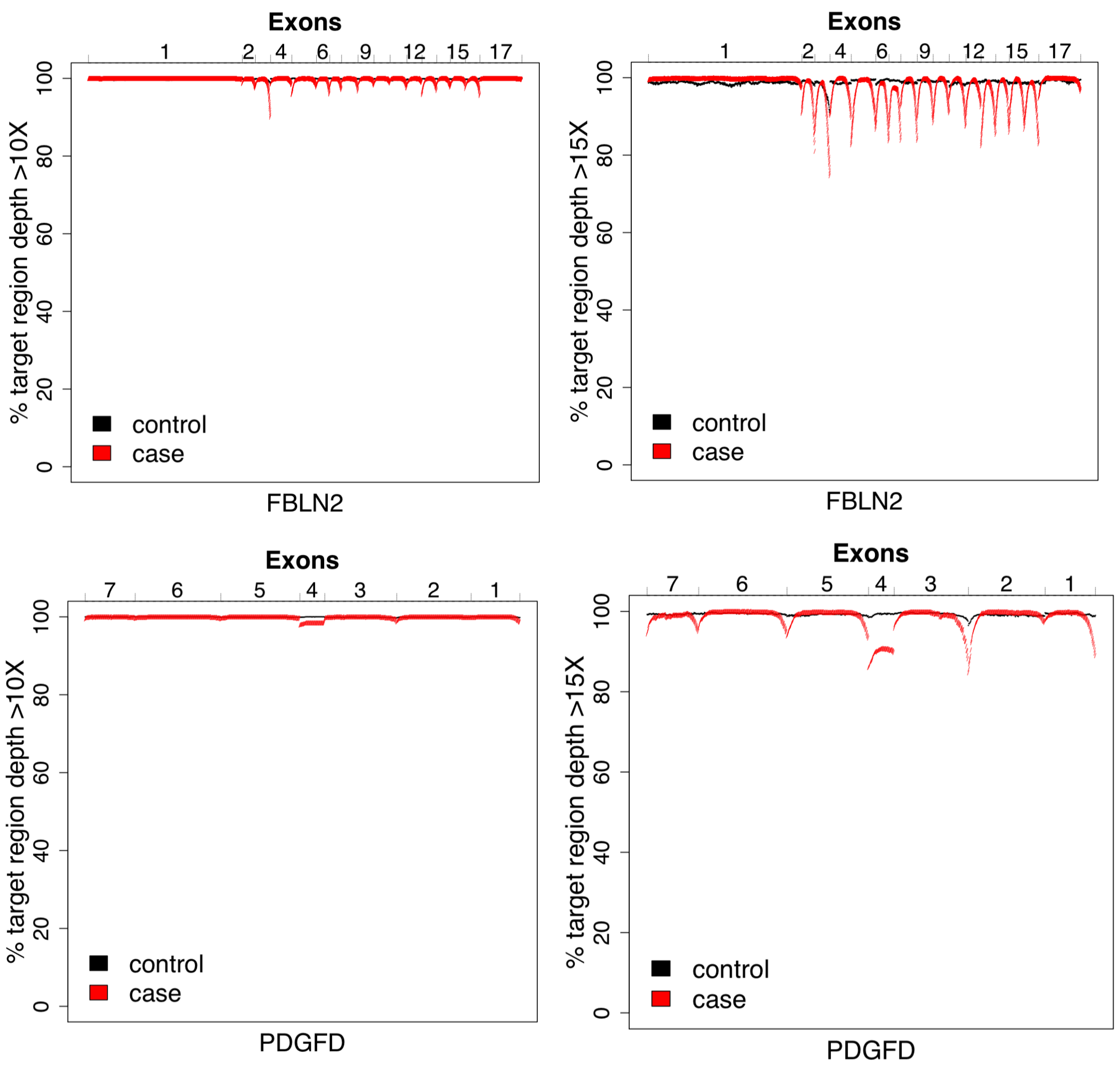
