## Supplemental Figure 4 for "Rare variant analysis of 4,241 pulmonary arterial hypertension cases from an international consortium implicate *FBLN2*, *PDGFD* and rare *de novo* variants in PAH"

**Supplementary Figure 4. Gene-based association analysis using 988 European APAH cases and 18,819 European controls (11,101 SPARK parents, 7,718 NFE gnomAD v2.1.1).** Results of a binomial test confined to rare, likely gene damaging (LGD) and predicted deleterious missense (D-Mis) variants or D-Mis only variants in 20,000 protein-coding genes. No genes met the Bonferroni-corrected threshold for significance.

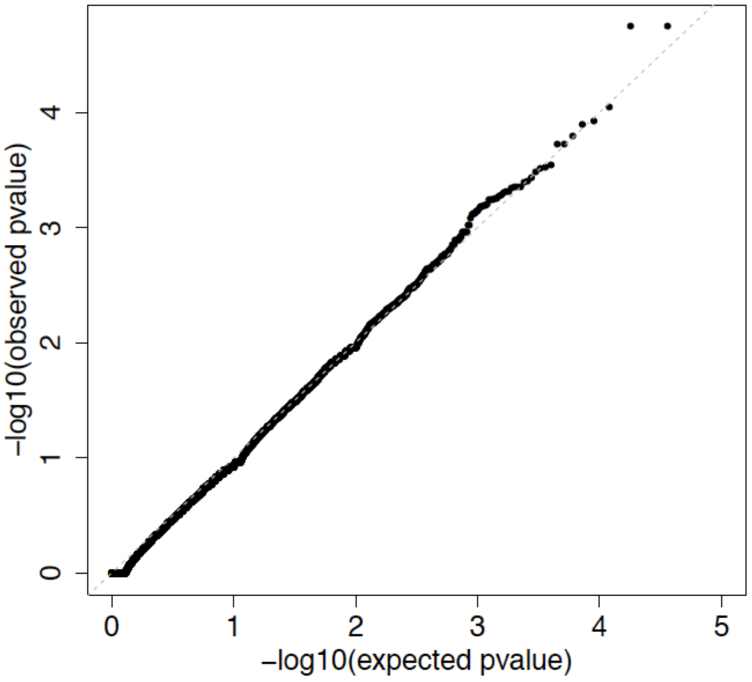
