## Supplemental Figure 5 for "Rare variant analysis of 4,241 pulmonary arterial hypertension cases from an international consortium implicate *FBLN2*, *PDGFD* and rare *de novo* variants in PAH"

**Supplementary Figure 5. Selection of single cell RNA-seq data available through the *Tabula Muris* Compendium. a)** Tissue, cell type and number of cells sequenced for data queried. Only tissues and cell types with >70 cells sequenced were included in the analyses. **b)** PCA analysis of tissue/cell type expression reveals a positive correlation between PC2 and endothelial cell type expression.

**a)**

| Tissue | Cell type | # cells sequenced |
|---|---|---|
| Lung | endothelial cell | 693 |
| Heart | endocardial cell | 165 |
| Aorta | endothelial cell | 188 |
| Heart | endothelial cell | 1177 |
| Heart | cardiac muscle cell | 133 |
| Aorta | erythrocyte | 91 |
| Lung | epithelial cell | 113 |
| Lung | classical monocyte | 90 |
| Lung | myeloid cell | 85 |
| Heart | leukocyte | 523 |
| Heart | myofibroblast cell | 178 |
| Lung | stromal cell | 423 |
| Heart | fibroblast | 2119 |
| Aorta | fibroblast | 70 |

**b)**

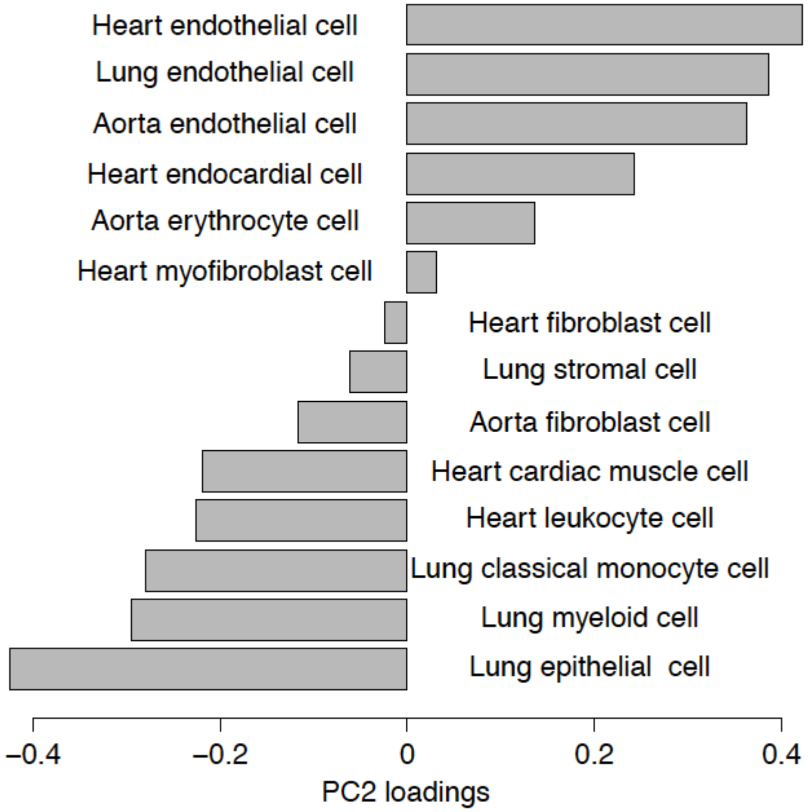
